# Paying upfront: successful initial infections protect against severe future infections

**DOI:** 10.1101/2025.09.24.676575

**Authors:** K.M. Talbott, A.E. Fleming-Davies, A.A. Perez-Umphrey, A.E. Henschen, J. Garrett-Larsen, F.E. Tillman, J.N. Weil, A.G. Arneson, S.J. Geary, E.R. Tulman, L.M. Childs, K.E. Langwig, D.M. Hawley, J.S. Adelman

## Abstract

Understanding the consistency with which individual hosts respond to repeated pathogen exposures is crucial for accurately modeling pathogen transmission and eco-evolutionary dynamics. When hosts face repeated pathogen exposures, immune memory is expected to reduce the probability and/or severity of subsequent infections, yet it remains unclear whether individuals remain consistent in their level of response relative to others. We investigated this question in house finches (*Haemorhous mexicanus*) from two populations varying in history of endemism of the bacterial pathogen *Mycoplasma gallisepticum* (MG). Individuals were confirmed to be MG-naive at capture and then experimentally inoculated twice with MG, allowing recovery between inoculations. We then asked if host responses to the second inoculation were predicted by responses to initial inoculation, sex, or population of origin. Our results suggest that individuals were not consistent in their relative response levels; rather, a successful initial infection provided protection against a severe second infection, increasing both tolerance and resistance. While we found no population differences in response to the second inoculation, males showed higher susceptibility to the second inoculation than females. Investigating and accounting for individual variation in response to subsequent exposures may improve the precision and accuracy of transmission models for wildlife pathogens.

## Introduction

At both individual and population levels, hosts show considerable heterogeneity in their responses to pathogens. One potentially critical, though often overlooked, driver of variation is prior exposure to a given pathogen [1,2]. While relevant empirical studies tend to focus on single pathogen exposures, hosts in a natural context often face repeated exposures to the same pathogen, as demonstrated by reinfection occurrence in humans and wildlife [3–8]. Thus, additional work is needed to understand how an individual’s initial response to pathogen exposure shapes or predicts subsequent responses to the same pathogen. Understanding the relative consistency of individual host responses is important not only for improving epidemiological models, but also for better understanding the physiological mechanisms and costs associated with different responses.

Host responses are often characterized in terms of susceptibility, resistance, and tolerance, all of which can vary within and between populations. Here, we define host susceptibility as the probability of infection following pathogen exposure, based on a binomial variable of “infected” or “uninfected” [2]; see Table 1. Resistance is defined as the ability of the host to reduce pathogen replication during infection, with more-resistant individuals showing lower pathogen loads than less-resistant conspecifics [9–11]. Tolerance is defined as the per-pathogen impact of infection on host pathology (a common proxy for host fitness), with more-tolerant hosts showing lower pathology at the same pathogen burden as less-tolerant hosts [11–15].

**Table 1.** Definitions of host responses to parasitism.

| Term | Definition |
| --- | --- |
| Susceptibility | Probability of infection following pathogen exposure at a given dose, based on a binomial variable of "infected" or "uninfected" |
| Resistance | Ability of the host to regulate within-host pathogen growth, quantified as pathogen load; individual hosts with higher resistance have relatively lower pathogen loads |
| Tolerance | Per-pathogen impact of infection on host fitness; here, more tolerant individuals experience less pathology per unit of pathogen than less tolerant individuals |

Quantifying heterogeneity in host responses in terms of susceptibility, tolerance, and resistance is key to refining predictions about epidemiological and host-pathogen coevolutionary dynamics. For example, higher resistance necessarily decreases an individual’s ability to infect another potential host (i.e., host competence; [16]). However, host tolerance may have distinct impacts on host competence, depending on the mode of pathogen transmission and effects of infection on host behavior [17–21]. Furthermore, tolerance and resistance may each have different ramifications for host population recovery following the emergence of novel pathogens [22], as well as for the evolution of pathogen virulence (i.e., harm caused to the host [23,24]).

In systems where repeated exposure to the same pathogen is common, the predicted relationship between individual responses to initial and subsequent exposures can depend on the type of host response examined, as well as the immune mechanisms involved. Vertebrate host responses to initial exposure depend largely on the innate immune response, a hallmark of which is inflammation [25,26]. In addition to directly killing invading pathogens, activated innate immune cells produce proinflammatory cytokines that help stimulate the subsequent development of an acquired immune response [27]. In turn, acquired immunity functions to reduce host susceptibility to a second exposure, or failing that, to help reduce pathogen burden during a second infection [25]. Because robust proinflammatory responses can incur damage to a host’s own tissues (e.g., immunopathology), muting inflammation has been proposed as a potential mechanism of tolerance [11,12,17,28,29]. However, it remains unclear whether this muting of inflammation during an initial infection reduces the development of acquired immunity, potentially increasing susceptibility or decreasing resistance in future pathogen exposures [30].

Despite the costs of immunopathology, theory predicts that selection may favor more pronounced inflammatory responses if they are associated with reduced susceptibility to a second infection, which we term the ‘Protective Infection Hypothesis’ [30]. Under the Protective Infection Hypothesis, we would predict that hosts susceptible to initial inoculation, especially those that become the sickest (i.e., low tolerance and/or resistance), would experience either decreased susceptibility to a second inoculation or less severe illness during second infection (i.e., high tolerance and/or resistance; see Figure 1). Alternatively, some individuals may be consistently less or more resistant, and/or less or more tolerant, relative to others. We term this the ‘Host Quality Hypothesis’. Under the Host Quality Hypothesis, we would predict that individuals who become infected and are the sickest (i.e., higher susceptibility, lower tolerance, and/or lower resistance) following initial inoculation would remain susceptible and become the sickest after a second inoculation. Thus, we expect tolerance or resistance during the first and second inoculations to be positively correlated under the Host Quality Hypothesis but negatively correlated under the Protective Infection Hypothesis (Figure 1).

**Figure 1.**
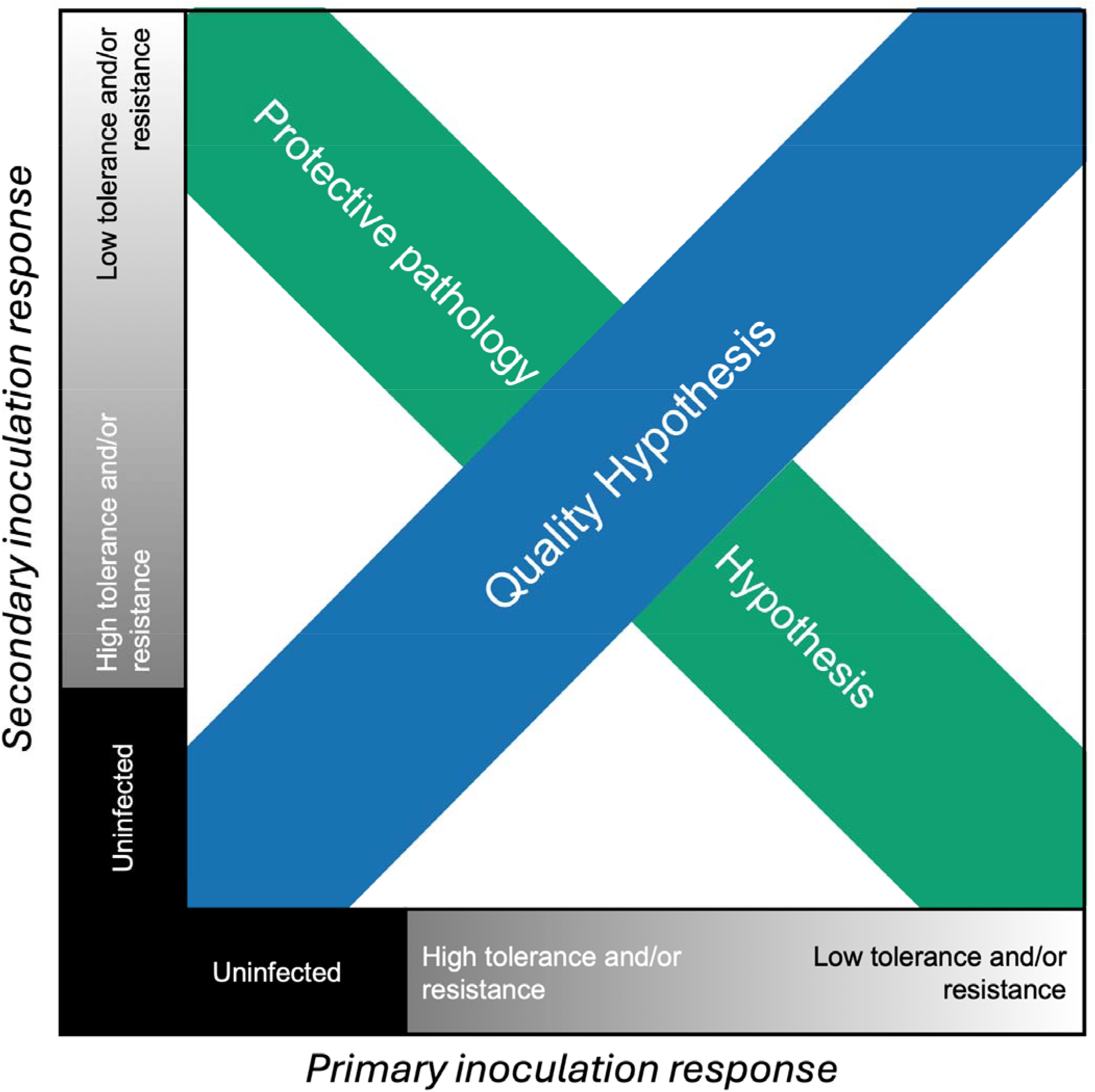
Hypothesized relationships between initial and second inoculation responses in individual house finches (*Haemorhous mexicanus*) inoculated with *Mycoplasma gallisepticum* (MG) bacteria twice. Note that responses form a gradient of infection severity and are not discrete response categories.

To test these hypotheses, we used house finches (*Haemorhous mexicanus*, ‘finches’) infected with the bacterium *Mycoplasma gallisepticum* (MG) as a model host-pathogen system. MG is a pathogen of poultry that spilled over into wild finches in eastern North America in the mid-1990’s and rapidly spread westward [31,32]. In finches, MG can cause severe conjunctivitis, rhinitis, and lethargy [33], which led to population declines immediately following spillover [32,34,35]. MG continues to circulate in populations where it has become endemic and experimental studies illustrate that individuals are susceptible to reinfection [1,36,37]. Recent work has also shown that MG exposure increases heterogeneity in susceptibility among individuals and reduces mean susceptibility to a second inoculation in finches from an MG-endemic population [2]. This indicates that while individuals gain some protection against a second infection, the extent of this protection varies among individuals. Here, we sampled finches from two distinct populations across the MG invasion gradient: Virginia (VA) and Arizona (AZ). MG has been endemic in the VA population for at least twenty years longer than the Tempe, AZ population sampled for this study [31,38], although MG was first detected in southern AZ in 2011 [39]. Therefore, the two study populations will be referred to as MG-endemic and MG-naive, respectively.

In past work, experimentally inoculated finches from MG-endemic populations showed higher tolerance [12,38,40], quicker recovery [40], and in some cases, lower MG burden (i.e., higher resistance [41], but see [12,38]) compared to finches from MG-naive populations. This suggests a history of stronger MG-mediated selection in MG-endemic populations. If this selection has favored robust immune memory over an effective initial response, we would predict a negative relationship between initial and second responses, as in the Protective Infection Hypothesis line in Figure 1; in this scenario, individuals from the MG-endemic population would show a stronger negative slope than individuals from the MG-naive population. Conversely, if non-memory responses have been selected for, we would expect a positive relationship between initial and second responses (i.e., individuals would fall along the Host Quality Hypothesis line of Figure 1, with birds from the MG-endemic population on the bottom left-hand side of the line and birds form the MG-naive population on the upper right-hand side).

If this selection has favored early (non-memory) responses over memory responses, we would expect populations to fall along the Host Quality Hypothesis line of Figure 1, with birds from the MG-endemic population generally better protected (bottom left-hand side) and birds form the MG-naive population less so (upper right-hand side). Conversely, if robust immune memory has been selected over initial (non-memory) responses, we would predict results more consistent with the Protective Infection Hypothesis line in Figure 1, but with birds from the MG-endemic population showing a more negative slope than individuals from the MG-naive population.

In addition to host population, host sex is another potential source of variation in the relationship between initial and second inoculation responses. For example, mortality was higher in adult male finches compared to adult females following initial MG emergence [42]. This could have resulted from sex differences in immune responses to MG or from sex-biased costs of similar immune responses. Either of these possibilities could lead to sex differences in MG responses, as seen in MG-inoculated canaries [43]. Similarly, sex differences could occur under either the Protective Infection Hypothesis or the Host Quality Hypothesis; our work here will help discern whether these competing hypotheses play a role in sex differences in MG response.

We tested these predictions and questions using an experimental approach in wild-caught finches of both sexes, captured from two populations that vary in MG endemism. All finches were given a standard priming dose (‘initial inoculation’) or carrier control, allowed to recover, and then inoculated a second time with one of four possible doses. We used a model selection approach to evaluate the extent to which initial inoculation responses, host sex, and host population of origin predicted susceptibility, resistance, and tolerance to a second inoculation at the individual level.

## Methods and Materials

### Experimental inoculations and data collection

We captured wild house finches in Tempe, AZ and Blacksburg, VA during 2021 and 2022; see Supplement for full capture and care methods. On days 3, 7, and 14 after arriving in captivity, birds were inspected for clinical signs of infection with MG (see eye scoring description above). On day 14, we collected blood samples to test for the presence of anti-MG antibodies, using a commercially available kit (99-09298, IDEXX, Westbrook, Maine) and previously published methods [44]. Only animals that showed no clinical signs before the experiments and no evidence of anti-MG antibodies (< 0.061 optical density at 630nm) were included in the study.

The experiment included two inoculations with *Mycoplasma gallisepticum* strain VA1994 (NCSU ADRL 7994-1), which was isolated soon after the emergence of MG in wild finches [31]. Finches were inoculated with an initial dose of 750 color changing units (CCU)/mL of MG and one of four doses (30, 100, 300, or 7,000 CCU/mL) during the second inoculation, using methods described previously [2]. The data from individuals in the MG-endemic population used in this analysis constitutes a subset of that collected for an earlier project focused on quantifying the impact of prior MG exposure on heterogeneity in susceptibility [2]. However, this work did not include the MG-naive population and focused solely on quantifying population-level variation in susceptibility, rather than the individual-level patterns considered here. See Figure 2 for a general experimental timeline and Table S1 for sample sizes by second MG dose, host population, and host sex.

**Figure 2.**
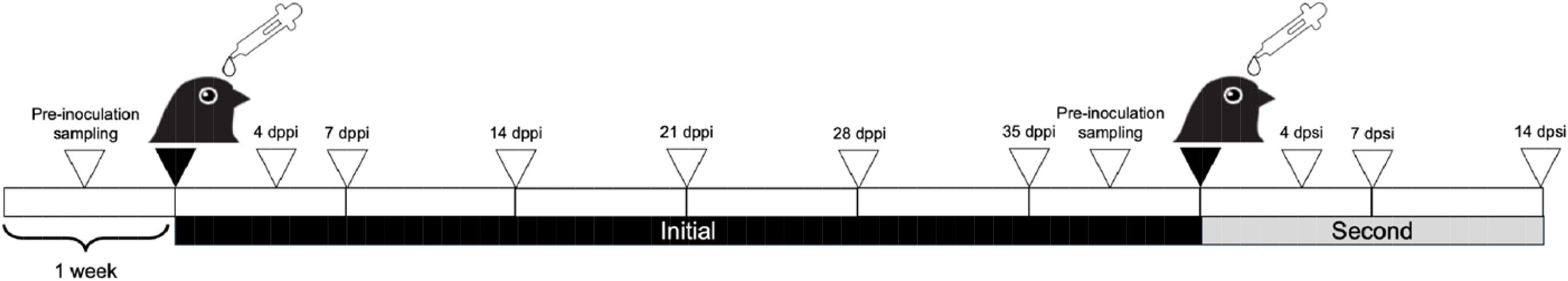
Experimental inoculation and sampling timeline of house finches (*Haemorhous mexicanus*) inoculated with *Mycoplasma gallisepticum* (‘MG’). White carrots indicate sampling dates and black carrots indicate inoculation dates (dpii = days post-initial inoculation, dpsi = days post-second inoculation). All finches were inoculated with an initial dose of 750 CCU/mL (color-changing units) of MG and one of four second doses (30, 100, 300, or 7,000). Sampling included visual scoring of eye pathology and/or swabs for assessing MG load. See Methods and Materials for additional details.

During both years (2021 and 2022), the second inoculation took place 42 days after the initial inoculation, by which time most birds had recovered; those that did not were removed from analysis (see ‘Statistical Analysis’ section). We used a visual scoring system to assess conjunctival pathology (‘eye scores’; see timeline below) [36]; briefly, each eye was assessed on a scale from 0 (no visible pathology) to 3 (eye nearly swollen shut), and scores from both eyes were summed. We then collected conjunctival swabs to quantify MG load. The conjunctiva of each eye was swabbed using a sterile cotton swab wetted with sterile tryptose phosphate broth (TPB) for 5 s and then wrung out in a tube of sterile TPB (300 µL). Samples were stored at −20°C until DNA extraction. MG genomic DNA was extracted from each broth sample by using Qiagen DNeasy 96 Blood and Tissue Kits (Qiagen, Valencia, CA), using methods described previously [45]. We quantified MG load from extracted DNA by using a qPCR assay with primers targeting MG’s *mgc2* gene [44] and methods as described previously [45]. All load values are transformed by log10(load + 1) unless otherwise noted.

We collected conjunctival eye swabs within 7 days prior to initial inoculation, and 7, 14 (2021 only), 40 or 41, 46, 49, 56, and 63 (2021 only) days following initial inoculation. Eye scores were collected 0, 7, 14, 21, 28, 35, 40 or 41, 46, and 56 days post-initial inoculation. These date ranges were chosen based on an expected peak in MG pathology and load at seven days following inoculation [12], with most birds recovered by 41 days following inoculation [2].

### Quantifying inoculation responses

Susceptibility to both initial and second inoculation were binomial variables reflecting whether an individual was successfully infected following MG inoculation. Finches with an eye score > 0 on 7, 14, 21, 28, 35, and/or 41 dpii, and/or a pathogen load > 15 copies on any of these dates, were considered successfully infected following initial inoculation; finches not meeting at least one of these criteria were scored as uninfected. Similarly, finches with an eye score > 0 on day 4, 7, 14, and/or 21 dpsi, and/or a load > 15 copies on any of these dates, were considered infected following the second inoculation; finches not meeting one of these criteria were scored as uninfected.

Because resistance refers to the ability of an individual to control pathogen replication, we used qPCR-derived pathogen loads as a proxy of resistance. Thus, individuals with relatively lower pathogen loads were classified as more resistant than those with relatively higher pathogen loads. Specifically, we quantified resistance as 7 dpi (days post-inoculation) MG loads following both initial and second inoculation (‘initial load’ and ‘second load’, respectively). This date was chosen because finches sampled in 2022 were missing load data from 14 dppi, and we wished to standardize the number of days post-inoculation for assessing initial and secondary resistance.

To estimate tolerance to initial and second inoculation, we used eye pathology (i.e., eye score) as a proxy of survival fitness, as higher eye scores are associated with a reduced probability of survival in wild finches [35]. We defined higher tolerance as lower eye scores per unit pathogen load, and estimated this with the negative residuals of a binomial model of eye score on pathogen load, such that larger values indicate higher tolerance. Because combined eye scores are a categorical variable between 0 and 6, we divided the maximum eye score recorded over the first two weeks following inoculation by 6, to give a variable between 0 and 1, following the approach of [38] for estimating tolerance. We then used a binomial generalized linear model to predict the maximum eye score recorded over the first two weeks following inoculation (divided by 6), using 7 dpi MG loads as a predictor (e.g., 7 dppi loads to calculate initial tolerance, and 7 dpsi loads to calculate second-inoculation tolerance). Note that hereafter, ‘maximum eye score’ refers to the maximum eye score recorded over the two weeks following a particular inoculation.

### Statistical analysis

We investigated factors predicting house finch susceptibility, tolerance, and resistance to the second inoculation with MG through a model selection approach using Akaike Information Criterion corrected for small sample sizes (AICc; [46]). Similar analyses for initial inoculation responses are available in the Supplement. Finches that showed signs of infection immediately before the second inoculation (eye score > 0 and/or MG load > 15 copies on 41 or 42 dpii) were excluded from analysis (n = 18). We also removed data from finches with incomplete records (n = 4), and controls that came up as qPCR positive at any point (n=14), likely due to low-level contamination of our qPCR assay [1]. For analyses of susceptibility to the second MG inoculation, we generated a list of candidate generalized linear models (GLMs) with a binomial error distribution and compared them by AICc by using the ‘aictab’ function in the *bbmle* package in R [47]. To investigate factors predicting resistance and tolerance to the second inoculation, we used a similar approach using linear models with a Gaussian error distribution. Candidate models included one or more of the following predictors: second MG dose, host population, host sex, and responses to the first (i.e., initial) MG inoculation. Responses to initial inoculation included initial susceptibility (i.e., infected or uninfected during the first MG challenge), initial MG load, maximum initial eye score, and initial tolerance residuals (initially infected finches only). The top models <2 ΔAICc for each analysis are reported. In cases where the null has the highest weight, no other models are reported. In cases where there are multiple models <2 ΔAICc and the top model (i.e., highest weighted) is also the simplest, only the top model is reported [48,49]. For top models, we assessed residuals using the *DHARMa* package [50]. All analyses were carried out in R version 2024.04.2+764 (R Core Team 2024).

Candidate models predicting susceptibility to the second inoculation (n = 84 finches) included the predictors listed above, as well as all possible interactions. Because we hypothesized distinct physiological processes in finches that successfully developed initial infection, compared to those that did not, we ran a follow-up model comparison using only finches that were successfully infected during initial inoculation (n = 37); predictors included those listed above apart from initial susceptibility.

To identify factors predicting resistance in finches successfully infected following the second inoculation (n = 22), we used the same predictors as described above. Within the MG-endemic population, relatively few birds (n=2/9) that were successfully infected from the first inoculation became reinfected after the second inoculation, reducing our ability to investigate consistency in resistance within this group. Therefore, we ran a second analysis in finches from the MG-naive population only (n = 13 finches that were successfully infected from first inoculation and became reinfected). Candidate models included the same predictors as listed above, apart from host population.

Finches that did not develop infection had no detectable eye pathology or MG load; therefore, these individuals would have similar tolerance residuals (i.e., very small) as successfully infected finches that had relatively low eye pathology scores. To avoid conflating these two scenarios, which likely reflect different physiological mechanisms, we investigated predictors of second-inoculation tolerance only in animals who developed infection (n=22). Candidate models included the same predictors as listed above. For the same reasons as outlined for analyses of resistance, we conducted a second analysis restricted to finches from the MG-naive population (n = 13). Similarly, candidate models included the same predictors listed above, apart from host population.

## Results

### Susceptibility to the second inoculation

Among all finches (n = 84), susceptibility to the second inoculation was positively correlated with the dose a finch received at second inoculation (Est = 3.90^-4^ ± 9.88^-5^ SE; Table S2), and was higher among male finches compared to females (Figure 3A; Est = 1.17 ± 0.69 SE). In addition, susceptibility to the second inoculation was lower for animals that had shown higher tolerance to initial infection (Est = −11.14 ± 5.39 SE); see Figure 3B and Table S2. Similar analyses for initial susceptibility are available in the Supplement. See Table S1 for sample sizes by second MG dose, host sex, and host population. In addition, we used a chi square test to confirm that susceptibility to the second inoculation was independent of susceptibility to the initial inoculation (X^2^_1_ = 4.45^-4^, p = 0.98).

**Figure 3.**
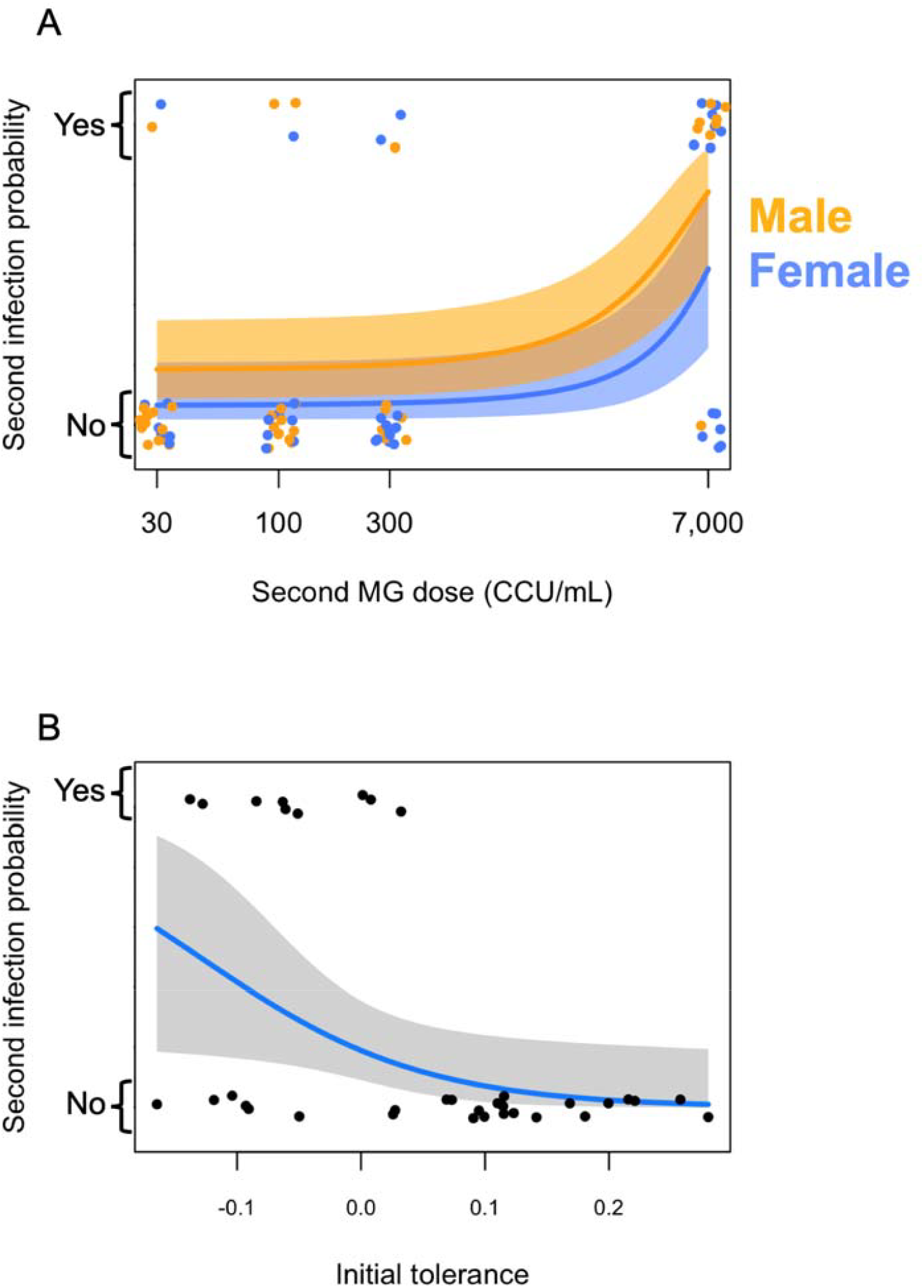
Shown are model predictions and 95% confidence intervals of a generalized linear model predicting the probability of infection following a second inoculation of *Mycoplasma gallisepticum* (MG) in house finches (*Haemorhous mexicanus*; n = 84). Panel A) infection probability at all second MG doses is higher for male (shown in yellow) than female finches (blue); note that the x-axis is log10 transformed but doses are not. Panel B) among finches successfully infected during their first MG exposure (n = 37), those with higher tolerance had a lower probability of infection following a second MG exposure. In both plots, individual points, representing data for individual finches, are jittered for easier viewing.

For initially infected finches (n = 37), second-inoculation susceptibility similarly increased with second MG dose (Est = 5.11^-4^ ± 2.50^-4^ SE), was higher for male finches compared to females (Est = 3.61 ± 1.89), and was lower for those with higher initial tolerance (Est = −12.04 ± 6.55). Susceptibility was also lower in MG-endemic finches compared to MG-naive finches (Est = −2.62 ± 1.39; see Table S3). See Table 2 for a summary of main results.

**Table 2.** Summary table of responses of house finches (*Haemorhous mexicanus*) to a second experimental inoculation with *Mycoplasma gallisepticum* bacteria (‘MG’). Responses (black column) are given in terms of susceptibility (S), resistance (R), and tolerance (T); see Table 1 for full definitions of each term. Potential predictors of each response type are listed in the top grey row. Those selected by model comparison as important predictors of each response are listed. Blank spaces indicate the factor was not selected as an important predictor. Results that are restricted to the subset of finches that were successfully infected following the first inoculation are marked with an asterisk*. Results restricted to finches from the MG-naive population (AZ) are marked with two asterisks**. Example interpretation, “Tolerance to a second inoculation is higher with higher second MG doses.”

| <u>Response to second inoculation</u> | Second MG dose | Host sex | Host population | Initial susceptibility | Initial resistance | Initial tolerance | Initial pathology |
| --- | --- | --- | --- | --- | --- | --- | --- |
| **Susceptibility**<br>(Probability of infection) | Higher S with higher second dose | Higher S in males than females | Higher S in MG-naïve than in MG-endemic* |  |  | Lower S with higher initial tolerance |  |
| **Resistance**<br>(Higher with lower MG load) | Lower R with higher second dose |  |  | Higher R in initial infected than in initial uninfected | Lower R with higher initial resistance** |  | Higher R with higher initial pathology** |
| **Tolerance**<br>(Higher with less pathology per unit pathogen load) | Higher T with higher second dose** |  |  | Higher T in initial infected than in initial uninfected** |  |  |  |

### Resistance to the second inoculation

Among finches that were successfully infected following the second inoculation (n = 22), second MG loads increased (i.e., resistance decreased) with higher second MG doses (Est = 1.73^-4^ ± 6.99^-5^ SE). In addition, initially uninfected finches had higher second loads (i.e., lower resistance) compared to initially infected finches (Est = 3.01 ± 0.47 SE); see Figure 4 and Table S4. Similar analyses for initial resistance are available in the Supplement.

**Figure 4.**
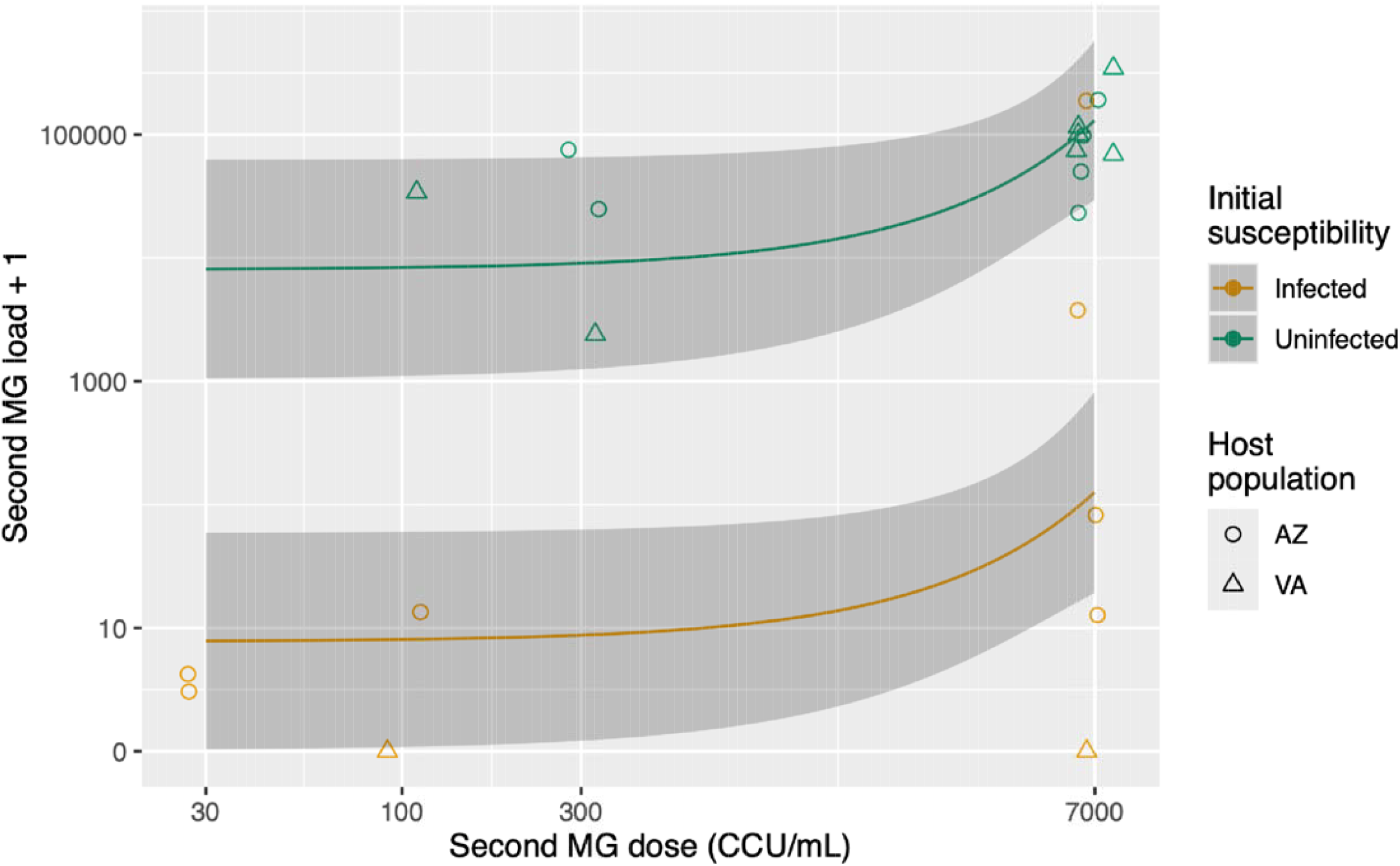
In house finches (*Haemorhous mexicanus*) inoculated sequentially with *Mycoplasma gallisepticum* (MG) bacteria, resistance (measured as MG load, log10-transformed on the y-axis) following the second inoculation is relatively higher in finches that became infected and developed relatively higher loads following the initial inoculation, and also varies with MG dose (x-axis). Shown are predictions and 95% confidence intervals from a linear model of resistance with second MG inoculum dose and initial susceptibility status as predictors (i.e., initially infected, in green or initially uninfected, in orange). Points represent data from individual finches (n = 22) and are jittered for ease of viewing. Although host population is not included in the model, points are shaped by host population (AZ in circles, VA in triangles). Note that the x-axis is log10 transformed for easier viewing, but doses are not transformed in the model.

Among finches from the MG-naive population (n = 13), initially uninfected finches had higher second loads (i.e., lower resistance) compared to initially infected finches (Est = 2.75 ± 0.74 SE). There was a negative relationship between initial and second MG loads (Est = −0.64 ± 0.17 SE). Similarly, there was a negative relationship between initial eye score and second MG loads (i.e., a positive relationship between initial eye pathology and second-inoculation resistance; Est = −1.09 ± 0.33). Note that these results reflect a top model set of three univariate models and that the modest sample size prohibited including a full model in the model comparison; see Table S5.

### Tolerance of the second inoculation

Among finches that became successfully infected following the second inoculation (n=22), none of our predictors (sex, host population, and responses to initial infection) improved predictions of second-inoculation tolerance compared to a null model. See Table S6 for details. Similar analyses for initial tolerance are available in the Supplement.

However, among MG-naive finches (n = 13), initially uninfected finches had lower second-inoculation tolerance than initially infected finches (Est = −0.31 ± 0.12). There was also a positive relationship between second MG dose and tolerance (Est = 4.02^-5^ ± 1.8^-5^); see Figure 5. Note that the null model was also included in the top model set (ΔAIC = 1.55, weight = 0.13); see Table S7.

**Figure 5.**
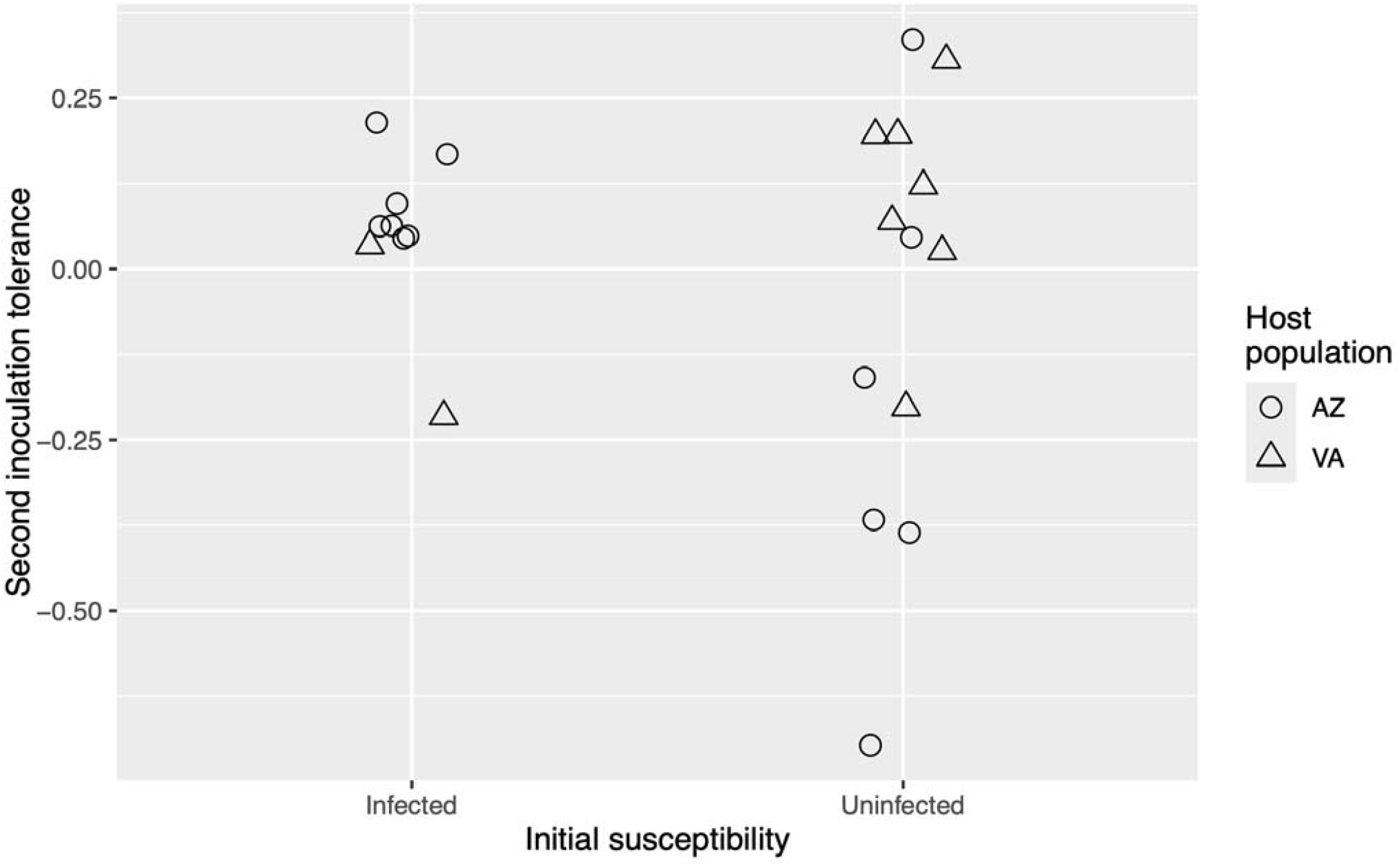
In house finches (*Haemorhous mexicanus*) from Arizona inoculated twice with *Mycoplasma gallisepticum* (MG) bacteria, tolerance following a second MG inoculation was higher in finches that were susceptible to initial inoculation (n = 7), compared to those not susceptible to initial inoculation (n = 6). Data are also shown from VA finches (triangles, n = 9), but these data were not included in the analysis of second inoculation tolerance. Tolerance is quantified as the residuals from a GLM predicting eye pathology from MG load; see Methods and Materials for additional details. Points represent tolerance values for individual finches.

## Discussion

Here we used an experimental approach in wild-caught house finches to investigate whether host responses to an initial inoculation with *Mycoplasma gallisepticum* (‘MG’) were predictive of responses to a second inoculation, and whether this relationship varied by host sex and/or population of origin. Specifically, we tested the competing hypotheses that individual host’s responses to the second inoculation were a) negatively associated with initial responses (‘Protective Infection Hypothesis’), or b) relatively consistent with their responses to initial inoculation (‘Host Quality Hypothesis’). If the Protective Infection Hypothesis were true, we would expect finches that developed infection and became the sickest would have a relatively lower probability of infection and/or would be less sick (i.e., higher resistance and/or tolerance) following the second inoculation (see Figure 1). Conversely, under the Host Quality Hypothesis, we would predict that individuals who became infected and were the sickest (i.e., low tolerance and/or resistance) following initial inoculation would also develop infection and become the sickest after the second inoculation.

We found moderate support for the Protective Infection Hypothesis. Initially infected finches had lower second MG loads than initially uninfected finches, suggesting that susceptibility to initial inoculation increased second-inoculation resistance. Among finches from a MG-naive population (i.e., those from AZ), initial and second loads were negatively associated, indicating that AZ finches with the lowest initial resistance showed the highest resistance to the second inoculation. Similarly, initially infected VA finches had higher second-inoculation tolerance than initially uninfected finches from the same population. Together, these results support the Protective Infection Hypothesis prediction that finches with a more severe initial infection would develop stronger protection, for example through immune memory, and thus experience less severe second infections. Similarly, humans infected with the novel SARS-Cov-2 virus show a positive relationship between disease severity and the production of neutralizing antibodies [51].

One might predict that this strategy – developing a relatively severe initial infection to avoid a severe second infection – would be selectively favored when the risk of repeated exposure is high. However, finches with relatively higher initial tolerance showed reduced second-inoculation susceptibility, indicating that there are multiple strategies by which individuals may avoid severe future infections. Importantly, both strategies reflect a multi-step selection process, requiring that individuals survive a successful initial infection. This may help explain the persistence of individual variation in immunopathology within host populations where MG has become endemic [30]. Additional work is needed to determine whether these patterns are general across finch populations and MG strains, as well as other host-pathogen systems.

In comparison, we found relatively less support for the Host Quality Hypothesis. One piece of supporting evidence is that individuals that became infected following the initial inoculation and had relatively lower tolerance (i.e., “lower quality” hosts) had a higher probability of a second infection, compared to those that had relatively higher tolerance during initial infection. However, we did not observe individual consistency in susceptibility; the numbers of individuals that were consistently susceptible (double-infection group) or unsusceptible to MG inoculation (zero-infection group) were consistent with a null model in which probability of second infection was independent of probability of first infection and thus initial susceptibility was not predictive of second susceptibility. Rather, the existence of the zero-infection and double-infection groups is better explained by variation in second MG doses, as the probability of infection following the second inoculation increased with higher second MG doses, which is an expected consequence of such increased pathogen pressure [1]. Similarly, we found no support for individual consistency in resistance or tolerance, leading us to the conclusion that the Protective Infection Hypothesis is better supported in this system than the Host Quality Hypothesis. This is contrary to recent work that has demonstrated individual consistency in immune profiles of wild mammalian hosts [52,53]. For example, in Soay sheep (*Ovis aries*), interleukin-4 and interferon-gamma showed longitudinal consistency, and were also positively associated with resistance to gastrointestinal helminths and coccidia, respectively [53]. Therefore, the probability of individual consistency in host responses to parasitism might vary by host taxon or host-pathogen dynamics (i.e., sterilizing immunity vs. chronic infection). Additional work is needed to address these possibilities.

It is also important to note that relatively few finches were susceptible to both inoculations, reducing our power to detect individual consistency in tolerance. Therefore, it remains unclear whether tolerance is a repeatable trait at the individual level, and further work is needed to address this question. In addition, we only quantified tolerance in successfully infected finches, rather than assigning high tolerance values to uninfected finches. We adopted this approach because distinct physiological responses could yield similar tolerance values when uninfected individuals are included in analysis. For example, because tolerance scores are quantified as a residual of a model predicting eye score from load, an uninfected individual with load and eye score of zero would have a similar score as any infected individual with load and pathology scores along the reaction norm. Therefore, to distinguish these processes, we recommend that researchers analyze infected and uninfected groups separately in studies that address variation in host tolerance at the individual level.

### Responses to second MG inoculation by host population and sex

Demographic variation in host responses to initial pathogen exposure are well documented. For example, host populations may evolve tolerance, resistance, and/or reduced susceptibility to emerging pathogens [38,41,54–58], resulting in inter-population variation in host responses based on pathogen endemism. For example, prior work in the house finch-MG system showed higher tolerance and resistance to MG in finches from MG-endemic versus MG-naive populations [12,38,40]. Therefore, we predicted that the Protective Infection Hypothesis was more likely to be supported in the MG-endemic population (VA), compared to the MG-naive population (AZ). Conversely, if the Host Quality Hypothesis were supported, we predicted that finches from the MG-endemic population would show higher resistance and tolerance following a second inoculation, compared to finches from the MG-naive population.

Neither of these hypotheses were directly supported, as population was not selected as an important predictor of tolerance or resistance to a second MG inoculation. It should be noted that relatively few MG-endemic finches were susceptible to the second inoculation, reducing our ability to detect population differences in second-inoculation tolerance. That said, the probability of a second infection was higher in finches with relatively lower initial tolerance, and the MG-naive population had relatively lower initial tolerance than the MG-endemic population. Indeed, an analysis restricted to initially infected finches showed that finches from the MG-naive population had a higher probability of infection following the second inoculation, relative to MG-endemic finches. Together, these results may hint at a subtle signal of selection - that the MG-endemic population contains a greater proportion of finches that show relatively high tolerance of an initial MG inoculation and reduced susceptibility to a second MG inoculation. Indeed, reduced susceptibility to subsequent infection may be an underappreciated benefit of tolerance in other wild vertebrate populations where tolerance to novel pathogens has evolved.

Within populations, host responses may also vary by sex. Male vertebrates are generally predicted to have higher susceptibility and lower resistance to initial pathogen exposure than females [59] (but see [60]); however, the role of host sex in predicting responses to multiple pathogen exposures is less clear. In the house finch-MG system, several studies have shown higher pathology, pathogen load, and infection duration in MG-inoculated males, compared to inoculated females [43,61–63]. However, other studies noted no sex differences in MG infection responses [36,64–66], or even a female bias in pathology [1,67]. Interestingly, we found that males were more susceptible to the second inoculation than females, although we found no effect of sex on initial susceptibility (see Supplement). This may indicate that females experience stronger protection from an initial infection under the Protective Infection Hypothesis. Stronger protection could be mediated by a female bias in immune memory, despite prior work suggesting a male bias in MG-specific antibody production in MG-inoculated songbirds [43,68]. Future work should address possible sex differences in antibody persistence and localization.

The male bias in second-inoculation susceptibility may have important implications for transmission. Male-driven pathogen transmission is well demonstrated in mammals [69–72], although avian systems have received relatively little attention beyond sexually transmitted infections [73,74]. Previous experimental work demonstrated faster MG transmission among male songbirds than females after a single MG inoculation [43,75]. Male finches also prefer to feed near MG-infected, rather than healthy, conspecifics, while females show no preference [76]; this behavior creates an opportunity for male-biased transmission at bird feeders, where MG transmission is known to occur [63,75,77]. Together, these studies and our results suggest that males could disproportionately drive MG transmission in wild finches, with a potential for complex interactions with exposure history.

Beyond susceptibility, we found no sex differences in second-inoculation responses (i.e., tolerance or resistance). Similarly, we found no sex differences in responses to initial inoculation; together, these results argue against the Host Quality Hypothesis in terms of sex. Interestingly, we noted lower initial loads in MG-endemic females compared to MG-naive females (see Supplement), indicating relatively higher initial resistance in MG-endemic females. This sex by population interaction in individuals with no prior exposure, who have no acquired immunity, could reflect sex-specific selective pressure of MG on finch innate immunity. Indeed, female-biased resistance is commonly documented in vertebrates [59], although the generality of this pattern in birds has been challenged [60,78,79]. More work is needed to determine whether there are sex-specific physiological benefits of high initial resistance to MG in MG-endemic populations. Given the relatively higher second-inoculation susceptibility of males (Fig 3A), it is possible that initial resistance is more adaptive in females than males; because females are less likely to experience second infections, limiting pathogen replication during the initial infection may be advantageous. If this hypothesis is borne out, the finch-MG system would comprise a rare opportunity to study sex-specific selective pressures of emerging pathogens on host immune function in wild vertebrates.

### Broader implications for epidemiology and host-pathogen coevolution

Our findings have important implications for transmission and host adaptation to emerging pathogens. We found greater support for the Protective Infection Hypothesis than the Host Quality Hypothesis, which broadly suggests that individual hosts are not consistent in their relative levels of response to repeated pathogen exposure. This indicates possible shifts in individual hosts’ contribution to pathogen transmission (i.e., competence). Relative to finches that fully resisted an initial inoculation, initially infected finches had lower second MG loads (i.e., higher resistance) and, in the case of birds from the MG-naive population, less pathology per unit pathogen load (i.e., higher tolerance) following their second inoculation. These shifts could facilitate individual movement from high-competence to low-competence, as the probability of MG transmission increases with pathogen load, regardless of prior exposure [80], and is negatively associated with pathology [43,81,82]. Thus, initially infected finches may play a relatively minor role in MG transmission during subsequent exposures, perhaps even helping to mute transmission; for example, recent work shows reduced transmission by MG-inoculated finches with higher tolerance (as measured by severity of pathology [20,21]). Conversely, initially uninfected finches may play a relatively larger role in MG transmission during subsequent exposures, as they showed relatively larger second-inoculation loads. Accounting for host role reversal across pathogen exposures may help improve the accuracy of epidemic models in this and other host-pathogen systems.

Our results also have implications for host adaptation to emerging pathogens. We suggest that finches face alternative strategies following an initial MG exposure: a) develop an active infection and reduce the severity of a future infection or b) fully resist an initial infection and risk developing a severe future infection. The relative benefits of each strategy could vary by several factors, including the probability of future MG exposures, the availability of resources needed for reducing pathogen replication and repairing tissues damaged through immunopathology, and competing physiological demands on available resources (e.g., reproduction, thermal regulation, coinfections, etc), and the durability of the protective benefit of infection. Although MG prevalence peaks seasonally, it is present year-round in MG-endemic finch populations [67]; thus, we predict that the probability of repeated MG exposure is moderate to high for individual finches. Therefore, selection may favor individuals that are susceptible to initial MG exposure, compared to those that are not, because of the benefit of reduced infection severity following future pathogen exposure. Furthermore, the benefit of this strategy may vary by sex. Additional work in this system and others is needed to understand sex-specific costs and benefits of different responses to initial pathogen exposure, especially during different seasons, when hosts face variation in pathogen pressure, as well as competing demands on physiological resources (e.g., immune function and reproduction).

## Ethics statement

Capture and collection in 2021 were approved by the United States Fish and Wildlife Service (MB158404, MB82600B). Capture in VA was approved by the Virginia Department of Game and Inland Fisheries (066646). Capture in AZ was approved by the State of Arizona Game and Fish Department (SP406779, SP841723). Birds were imported to Tennessee through Tennessee Scientific Collecting Permit 2252 and Importation Permits 35257117 and 36586049. All capture, handling, and care procedures were approved by the Virginia Tech (VT) Institutional Animal Care and Use Committee (IACUC), Arizona State University IACUC, and University of Memphis IACUC prior to the start of work.

## Authors’ contributions

K.M.T.: conceptualization, methodology, software, formal analysis, data curation, writing (original draft, review & editing, visualization; A.E.F.: conceptualization, methodology, software, formal analysis, data curation, writing (review & editing), supervision, project administration, funding acquisition; A.A.P.: investigation, data curation, writing (review & editing); A.E.H.: investigation, data curation, writing (review & editing); J.G.: investigation and writing (review & editing); F.E.T.: investigation and writing (review & editing); J.N.W.: investigation, writing (review & editing); A.G.A.: investigation; S.J.G.: conceptualization, methodology, project administration, funding acquisition; E.R.T.: conceptualization, methodology, project administration, funding acquisition; L.M.C.: conceptualization, methodology, software, writing (review & editing), project administration, funding acquisition; K.E.L.: conceptualization, methodology, software, writing (review & editing), project administration, funding acquisition; D.M.H.: conceptualization, methodology, investigation, resources, writing (review & editing), supervision, project administration, funding acquisition; J.S.A.: conceptualization, methodology, formal analysis, investigation, resources, data curation, writing (review & editing), supervision, project administration, funding acquisition.

## Data accessibility

Data and code are deposited on https://github.com/talbottkm/mg_response_pop_sex/.

## Acknowledgment

We thank John Brule, Annabel Coyle, Noelle Hodges, Marissa Langager, Ama Owusu-Attakorah, Sara Teemer, Caro Vela, and Chava Weitzman for assistance with lab and field data collection. We also thank Kevin McGraw for assistance in Arizona finch capture.

## Declaration of AI use

We have not used AI-assisted technologies in creating this article.

## References

1. Leon AE, Hawley DM. 2017 Host Responses to Pathogen Priming in a Natural Songbird Host. EcoHealth 14, 793–804. (doi:10.1007/s10393-017-1261-x)

2. Hawley DM, Pérez-Umphrey AA, Adelman JS, Fleming-Davies AE, Garrett-Larsen J, Geary SJ, Childs LM, Langwig KE. 2024 Prior exposure to pathogens augments host heterogeneity in susceptibility and has key epidemiological consequences. PLOS Pathog. 20, e1012092. (doi:10.1371/journal.ppat.1012092)

3. Frank AL, Taber LH. 1983 Variation in frequency of natural reinfection with influenza A viruses. J. Med. Virol. 12, 17–23. (doi:10.1002/jmv.1890120103)

4. Trottier H, Ferreira S, Thomann P, Costa MC, Sobrinho JS, Prado JCM, Rohan TE, Villa LL, Franco EL. 2010 Human Papillomavirus Infection and Reinfection in Adult Women: the Role of Sexual Activity and Natural Immunity. Cancer Res. 70, 8569–8577. (doi:10.1158/0008-5472.CAN-10-0621)

5. Quinnell RJ, Pullan RL, Breitling LPh, Geiger SM, Cundill B, Correa‐Oliveira R, Brooker S, Bethony JM. 2010 Genetic and Household Determinants of Predisposition to Human Hookworm Infection in a Brazilian Community. J. Infect. Dis. 202, 954–961. (doi:10.1086/655813)

6. Lier T, Do DT, Johansen MV, Nguyen TH, Dalsgaard A, Asfeldt AM. 2014 High Reinfection Rate after Preventive Chemotherapy for Fishborne Zoonotic Trematodes in Vietnam. PLoS Negl. Trop. Dis. 8, e2958. (doi:10.1371/journal.pntd.0002958)

7. Adrielle Dos Santos L et al. 2021 Recurrent COVID-19 including evidence of reinfection and enhanced severity in thirty Brazilian healthcare workers. J. Infect. 82, 399–406. (doi:10.1016/j.jinf.2021.01.020)

8. Müller‐Klein N, Heistermann M, Strube C, Franz M, Schülke O, Ostner J. 2019 Exposure and susceptibility drive reinfection with gastrointestinal parasites in a social primate. Funct. Ecol. 33, 1088–1098. (doi:10.1111/1365-2435.13313)

9. Gandon S, Michalakis Y. 2000 Evolution of parasite virulence against qualitative or quantitative host resistance. Proc. R. Soc. Lond. B Biol. Sci. 267, 985–990. (doi:10.1098/rspb.2000.1100)

10. Råberg L, Sim D, Read AF. 2007 Disentangling Genetic Variation for Resistance and Tolerance to Infectious Diseases in Animals. Science 318, 812–814. (doi:10.1126/science.1148526)

11. Råberg L, Graham AL, Read AF. 2009 Decomposing health: tolerance and resistance to parasites in animals. Philos. Trans. R. Soc. B Biol. Sci. 364, 37–49. (doi:10.1098/rstb.2008.0184)

12. Adelman JS, Kirkpatrick L, Grodio JL, Hawley DM. 2013 House Finch Populations Differ in Early Inflammatory Signaling and Pathogen Tolerance at the Peak of Mycoplasma gallisepticum Infection. Am. Nat. 181, 674–689. (doi:10.1086/670024)

13. Caldwell RM, Schafer JF, Compton LE, Patterson FL. 1958 Tolerance to Cereal Leaf Rusts. Science 128, 714–715. (doi:10.1126/science.128.3326.714)

14. Kutzer MAM, Armitage SAO. 2016 Maximising fitness in the face of parasites: a review of host tolerance. Zoology 119, 281–289. (doi:10.1016/j.zool.2016.05.011)

15. Little TJ, Shuker DM, Colegrave N, Day T, Graham AL. 2010 The Coevolution of Virulence: Tolerance in Perspective. PLoS Pathog. 6, e1001006. (doi:10.1371/journal.ppat.1001006)

16. Martin LB et al. 2019 Extreme Competence: Keystone Hosts of Infections. Trends Ecol. Evol. 34, 303–314. (doi:10.1016/j.tree.2018.12.009)

17. Adelman JS, Hawley DM. 2017 Tolerance of infection: A role for animal behavior, potential immune mechanisms, and consequences for parasite transmission. Horm. Behav. 88, 79–86. (doi:10.1016/j.yhbeh.2016.10.013)

18. Lloyd-Smith JO, Schreiber SJ, Kopp PE, Getz WM. 2005 Superspreading and the effect of individual variation on disease emergence. Nature 438, 355–359. (doi:10.1038/nature04153)

19. Burgan SC, Gervasi SS, Johnson LR, Martin LB. 2019 How Individual Variation in Host Tolerance Affects Competence to Transmit Parasites. Physiol. Biochem. Zool. 92, 49–57. (doi:10.1086/701169)

20. Ruden RM, Adelman JS. 2021 Disease tolerance alters host competence in a wild songbird. Biol. Lett. 17, 20210362. (doi:10.1098/rsbl.2021.0362)

21. Henschen AE, Tillman FE, Ruston SC, Hawley DM, Adelman JS. 2025 Host Disease Tolerance Predicts Transmission Probability for a Songbird Pathogen. Ecol. Evol. 15, e70882. (doi:10.1002/ece3.70882)

22. Wilber MQ, DeMarchi JA, Briggs CJ, Streipert S. 2024 Rapid Evolution of Resistance and Tolerance Leads to Variable Host Recoveries following Disease-Induced Declines. Am. Nat., 000–000. (doi:10.1086/729437)

23. Pfennig KS. 2001 Evolution of pathogen virulence: the role of variation in host phenotype. Proc. R. Soc. Lond. B Biol. Sci. 268, 755–760. (doi:10.1098/rspb.2000.1582)

24. Miller MR, White A, Boots M. 2005 The evolution of host resistance: Tolerance and control as distinct strategies. J. Theor. Biol. 236, 198–207. (doi:10.1016/j.jtbi.2005.03.005)

25. Janeway CA. 2001 How the immune system protects the host from infection. Microbes Infect. 3, 1167–1171. (doi:10.1016/S1286-4579(01)01477-0)

26. Ashley NT, Weil ZM, Nelson RJ. 2012 Inflammation: Mechanisms, Costs, and Natural Variation. Annu. Rev. Ecol. Evol. Syst. 43, 385–406. (doi:10.1146/annurev-ecolsys-040212-092530)

27. Beutler B. 2004 Innate immunity: an overview. Mol. Immunol. 40, 845–859. (doi:10.1016/j.molimm.2003.10.005)

28. Sears BF, Rohr JR, Allen JE, Martin LB. 2011 The economy of inflammation: when is less more? Trends Parasitol. 27, 382–387. (doi:10.1016/j.pt.2011.05.004)

29. Medzhitov R, Schneider DS, Soares MP. 2012 Disease Tolerance as a Defense Strategy. Science 335, 936–941. (doi:10.1126/science.1214935)

30. Cressler CE, Adelman JS. 2024 Links between Innate and Adaptive Immunity Can Favor Evolutionary Persistence of Immunopathology. Integr. Comp. Biol., icae105. (doi:10.1093/icb/icae105)

31. Ley DH, Berkhoff JE, McLaren JM. 1996 Mycoplasma gallisepticum Isolated from House Finches (Carpodacus mexicanus) with Conjunctivitis. Avian Dis. 40, 480. (doi:10.2307/1592250)

32. Hochachka WM, Dhondt AA. 2000 Density-dependent decline of host abundance resulting from a new infectious disease. Proc. Natl. Acad. Sci. 97, 5303–5306. (doi:10.1073/pnas.080551197)

33. Kollias GV, Sydenstricker KV, Kollias HW, Ley DH, Hosseini PR, Connolly V, Dhondt AA. 2004 Experimental infection of house finches with Mycoplasma gallisepticum. J. Wildl. Dis. 40, 79–86. (doi:10.7589/0090-3558-40.1.79)

34. Dhondt AA, Tessaglia DL, Slothower RL. 1998 Epidemic mycoplasmal conjunctivitis in house finches from eastern North America. J. Wildl. Dis. 34, 265–280. (doi:10.7589/0090-3558-34.2.265)

35. Faustino CR, Jennelle CS, Connolly V, Davis AK, Swarthout EC, Dhondt AA, Cooch EG. 2004 Mycoplasma gallisepticum infection dynamics in a house finch population: seasonal variation in survival, encounter and transmission rate. J. Anim. Ecol. 73, 651–669. (doi:10.1111/j.0021-8790.2004.00840.x)

36. Sydenstricker KV, Dhondt AA, Ley DH, Kollias GV. 2005 Re-exposure of captive house finches that recovered from Mycoplasma gallisepticum infection. J. Wildl. Dis. 41, 326–333. (doi:10.7589/0090-3558-41.2.326)

37. Weitzman CL, Ceja G, Leon AE, Hawley DM. 2022 Protection Generated by Prior Exposure to Pathogens Depends on both Priming and Challenge Dose. Infect. Immun. 90. (doi:10.1128/iai.00537-21)

38. Henschen AE, Vinkler M, Langager MM, Rowley AA, Dalloul RA, Hawley DM, Adelman JS. 2023 Rapid adaptation to a novel pathogen through disease tolerance in a wild songbird. PLOS Pathog. 19, e1011408. (doi:10.1371/journal.ppat.1011408)

39. Staley M, Bonneaud C, McGraw KJ, Vleck CM, Hill GE. 2018 Detection of Mycoplasma gallisepticum in House Finches (Haemorhous mexicanus) from Arizona. Avian Dis. 62, 14–17. (doi:10.1637/11610-021317-Reg.1)

40. Bonneaud C, Tardy L, Giraudeau M, Hill GE, McGraw KJ, Wilson AJ. 2019 Evolution of both host resistance and tolerance to an emerging bacterial pathogen. Evol. Lett. 3, 544–554. (doi:10.1002/evl3.133)

41. Bonneaud C, Balenger SL, Russell AF, Zhang J, Hill GE, Edwards SV. 2011 Rapid evolution of disease resistance is accompanied by functional changes in gene expression in a wild bird. Proc. Natl. Acad. Sci. 108, 7866–7871. (doi:10.1073/pnas.1018580108)

42. Nolan PM, Hill GE, Stoehr AM. 1998 Sex, size, and plumage redness predict house finch survival in an epidemic. Proc. R. Soc. Lond. B Biol. Sci. 265, 961–965. (doi:10.1098/rspb.1998.0384)

43. Sauer EL, Connelly C, Perrine W, Love AC, DuRant SE. 2024 Male pathology regardless of behaviour drives transmission in an avian host–pathogen system. J. Anim. Ecol. 93, 36–44. (doi:10.1111/1365-2656.14026)

44. Grodio JL, Dhondt KV, O’Connell PH, Schat KA. 2008 Detection and quantification of Mycoplasma gallisepticum genome load in conjunctival samples of experimentally infected house finches (Carpodacus mexicanus) using real-time polymerase chain reaction. Avian Pathol. 37, 385–391. (doi:10.1080/03079450802216629)

45. Hawley DM, Osnas EE, Dobson AP, Hochachka WM, Ley DH, Dhondt AA. 2013 Parallel Patterns of Increased Virulence in a Recently Emerged Wildlife Pathogen. PLoS Biol. 11, e1001570. (doi:10.1371/journal.pbio.1001570)

46. Burnham KP, Anderson DR, editors. 2004 Model Selection and Multimodel Inference. New York, NY: Springer. (doi:10.1007/b97636)

47. Bolker B, Team RDC, Giné-Vázquez I. 2023 bbmle: Tools for General Maximum Likelihood Estimation.

48. Richards SA. 2008 Dealing with overdispersed count data in applied ecology. J. Appl. Ecol. 45, 218–227. (doi:10.1111/j.1365-2664.2007.01377.x)

49. Richards SA, Whittingham MJ, Stephens PA. 2011 Model selection and model averaging in behavioural ecology: the utility of the IT-AIC framework. Behav. Ecol. Sociobiol. 65, 77–89. (doi:10.1007/s00265-010-1035-8)

50. Hartig F, Lohse L, leite M de S. 2024 DHARMa: Residual Diagnostics for Hierarchical (Multi-Level /Mixed) Regression Models.

51. Garcia-Beltran WF et al. 2021 COVID-19-neutralizing antibodies predict disease severity and survival. Cell 184, 476-488.e11. (doi:10.1016/j.cell.2020.12.015)

52. Arriero E, Wanelik KM, Birtles RJ, Bradley JE, Jackson JA, Paterson S, Begon M. 2017 From the animal house to the field: Are there consistent individual differences in immunological profile in wild populations of field voles (Microtus agrestis)? PLOS ONE 12, e0183450. (doi:10.1371/journal.pone.0183450)

53. Corripio-Miyar Y et al. 2025 T-helper cell phenotypes are repeatable, positively correlated, and associated with helminth infection in wild Soay sheep. Discov. Immunol. 4, kyae017. (doi:10.1093/discim/kyae017)

54. Kerr PJ. 2012 Myxomatosis in Australia and Europe: A model for emerging infectious diseases. Antiviral Res. 93, 387–415. (doi:10.1016/j.antiviral.2012.01.009)

55. Atkinson CT, Saili KS, Utzurrum RB, Jarvi SI. 2013 Experimental Evidence for Evolved Tolerance to Avian Malaria in a Wild Population of Low Elevation Hawai’i ‘Amakihi (Hemignathus virens). EcoHealth 10, 366–375. (doi:10.1007/s10393-013-0899-2)

56. Savage AE, Zamudio KR. 2016 Adaptive tolerance to a pathogenic fungus drives major histocompatibility complex evolution in natural amphibian populations. Proc. R. Soc. B Biol. Sci. 283, 20153115. (doi:10.1098/rspb.2015.3115)

57. Langwig KE et al. 2017 Vaccine Effects on Heterogeneity in Susceptibility and Implications for Population Health Management. mBio 8, e00796–17. (doi:10.1128/mBio.00796-17)

58. Voyles J et al. 2018 Shifts in disease dynamics in a tropical amphibian assemblage are not due to pathogen attenuation. Science 359, 1517–1519. (doi:10.1126/science.aao4806)

59. Klein SL, Flanagan KL. 2016 Sex differences in immune responses. Nat. Rev. Immunol. 16, 626–638. (doi:10.1038/nri.2016.90)

60. Valdebenito JO, Halimubieke N, Lendvai ÁZ, Figuerola J, Eichhorn G, Székely T. 2021 Seasonal variation in sex-specific immunity in wild birds. Sci. Rep. 11, 1349. (doi:10.1038/s41598-020-80030-9)

61. Hawley DM, Jennelle CS, Sydenstricker KV, Dhondt AA. 2007 Pathogen resistance and immunocompetence covary with social status in house finches (Carpodacus mexicanus). Funct. Ecol. 21, 520–527. (doi:10.1111/j.1365-2435.2007.01254.x)

62. Love AC, Foltz SL, Adelman JS, Moore IT, Hawley DM. 2016 Changes in corticosterone concentrations and behavior during Mycoplasma gallisepticum infection in house finches (Haemorhous mexicanus). Gen. Comp. Endocrinol. 235, 70–77. (doi:10.1016/j.ygcen.2016.06.008)

63. Moyers SC, Adelman JS, Farine DR, Thomason CA, Hawley DM. 2018 Feeder density enhances house finch disease transmission in experimental epidemics. Philos. Trans. R. Soc. B Biol. Sci. 373, 20170090. (doi:10.1098/rstb.2017.0090)

64. Thomason CA, Leon A, Kirkpatrick LT, Belden LK, Hawley DM. 2017 Eye of the Finch: characterization of the ocular microbiome of house finches in relation to mycoplasmal conjunctivitis: Ocular microbiome and house finch conjunctivitis. Environ. Microbiol. 19, 1439–1449. (doi:10.1111/1462-2920.13625)

65. Vinkler M, Leon AE, Kirkpatrick L, Dalloul RA, Hawley DM. 2018 Differing House Finch Cytokine Expression Responses to Original and Evolved Isolates of Mycoplasma gallisepticum. Front. Immunol. 9.

66. Weitzman CL, Rostama B, Thomason CA, May M, Belden LK, Hawley DM. 2021 Experimental test of microbiome protection across pathogen doses reveals importance of resident microbiome composition. FEMS Microbiol. Ecol. 97, fiab141. (doi:10.1093/femsec/fiab141)

67. Altizer S, Davis AK, Cook KC, Cherry JJ. 2004 Age, sex, and season affect the risk of mycoplasmal conjunctivitis in a southeastern house finch population. Can. J. Zool. 82, 755–763. (doi:10.1139/z04-050)

68. Gates DE, Staley M, Tardy L, Giraudeau M, Hill GE, McGraw KJ, Bonneaud C. 2021 Levels of pathogen virulence and host resistance both shape the antibody response to an emerging bacterial disease. Sci. Rep. 11, 8209. (doi:10.1038/s41598-021-87464-9)

69. Ferrari N, Cattadori IM, Nespereira J, Rizzoli A, Hudson PJ. 2003 The role of host sex in parasite dynamics: field experiments on the yellow-necked mouse Apodemus flavicollis: Role of host sex in parasite dynamics. Ecol. Lett. 7, 88–94. (doi:10.1046/j.1461-0248.2003.00552.x)

70. Grear DA, Perkins SE, Hudson PJ. 2009 Does elevated testosterone result in increased exposure and transmission of parasites? Ecol. Lett. 12, 528–537. (doi:10.1111/j.1461-0248.2009.01306.x)

71. Leu ST, Sah P, Krzyszczyk E, Jacoby A-M, Mann J, Bansal S. 2020 Sex, synchrony, and skin contact: integrating multiple behaviors to assess pathogen transmission risk. Behav. Ecol. 31, 651–660. (doi:10.1093/beheco/araa002)

72. VanderWaal KL, Atwill ER, Hooper S, Buckle K, McCowan B. 2013 Network structure and prevalence of Cryptosporidium in Belding’s ground squirrels. Behav. Ecol. Sociobiol. 67, 1951–1959. (doi:10.1007/s00265-013-1602-x)

73. Kulkarni S, Heeb P. 2007 Social and sexual behaviours aid transmission of bacteria in birds. Behav. Processes 74, 88–92. (doi:10.1016/j.beproc.2006.10.005)

74. Sheldon BC. 1993 Sexually transmitted disease in birds: occurrence and evolutionary significance. Philos. Trans. R. Soc. Lond. B. Biol. Sci. 339, 491–497. (doi:10.1098/rstb.1993.0044)

75. Adelman JS, Moyers SC, Farine DR, Hawley DM. 2015 Feeder use predicts both acquisition and transmission of a contagious pathogen in a North American songbird. Proc. R. Soc. B Biol. Sci. 282, 20151429. (doi:10.1098/rspb.2015.1429)

76. Bouwman KM, Hawley DM. 2010 Sickness behaviour acting as an evolutionary trap? Male house finches preferentially feed near diseased conspecifics. Biol. Lett. 6, 462–465. (doi:10.1098/rsbl.2010.0020)

77. Dhondt AA, Dhondt KV, Hawley DM, Jennelle CS. 2007 Experimental evidence for transmission of Mycoplasma gallisepticum in house finches by fomites. Avian Pathol. 36, 205–208. (doi:10.1080/03079450701286277)

78. Kelly CD, Stoehr AM, Nunn C, Smyth KN, Prokop ZM. 2018 Sexual dimorphism in immunity across animals: a meta‐analysis. Ecol. Lett. 21, 1885–1894. (doi:10.1111/ele.13164)

79. Valdebenito JO, Jones W, Székely T. 2024 Evolutionary drivers of sex-specific parasite prevalence in wild birds. Proc. R. Soc. B Biol. Sci. 291, 20241013. (doi:10.1098/rspb.2024.1013)

80. Leon AE, Fleming-Davies AE, Adelman JS, Hawley DM. 2025 Pathogen priming alters host transmission potential and predictors of transmissibility in a wild songbird species. mSphere, e00886-24. (doi:10.1128/msphere.00886-24)

81. Adelman JS, Carter AW, Hopkins WA, Hawley DM. 2013 Deposition of pathogenic Mycoplasma gallisepticum onto bird feeders: host pathology is more important than temperature-driven increases in food intake. Anim. Behav. 9, 20130594. (doi:10.1098/rsbl.2013.0594)

82. Bonneaud C, Tardy L, Hill GE, McGraw KJ, Wilson AJ, Giraudeau M. 2020 Experimental evidence for stabilizing selection on virulence in a bacterial pathogen. Evol. Lett. 4, 491–501. (doi:10.1002/evl3.203)

