## Supplementary Materials for "Paying upfront: successful initial infections protect against severe future infections"

### Supplement

#### Finch capture and care

From June - August 2021, juvenile house finches were captured in and near Blacksburg, Virginia (VA) and Tempe, Arizona USA (AZ) using baited feeder traps and mist nets. Finches showing signs of conjunctivitis were released at the site of capture; all others were fitted with a metal, uniquely-numbered leg band for identification. Finches captured in VA were transported to Virginia Tech via automobile using approved transport methods; finches captured in AZ were transported to Arizona State University in similar fashion. Finches from AZ were held individually in cages (60 × 40 × 30 cm) in the Arizona State University vivarium for 2-8d, depending upon day of capture. These animals were transported via commercial aircraft to the University of Memphis in International Air Transport Association (IATA)-approved pet carriers that had been modified to hold birds. Briefly, each carrier (67 x 46 x 52 cm) included five perches, absorbent material on the floor, and sufficient seed and fruit for finches to remain satiated and hydrated during the journey. Airline regulations require that no water be present in the cage, which is why we substituted fruit. Each carrier housed a maximum of 20 animals and the total time in transit was approximately 20 hours.

We also captured finches in Tempe, AZ in 2022, using the same procedures as above. However, because of airline cancellations, we transported finches to the University of Memphis via automobile in the same carriers as above, with water and seed bowls added to the carriers. This journey took approximately 22 hours and animals were inspected visually for any signs of distress at least every 4 hours. No animals died during transit in either year.

Upon arrival at the University of Memphis or Virginia Tech, animals were housed either individually or in pairs in cages (76 x 46 x 46 cm) with *ad libitum* access to water, grit, and food (a 20:80 blend of black oil sunflower seed:Roudybush Maintenance Nibbles; Roudybush, Inc., Woodland, CA). Rooms were maintained at a lighting regime of 12 hrs light: 12 hrs dark and constant temperature of approximately 22°C. Animals were treated prophylactically to minimize impacts of common parasites found in wild birds. To reduce risk of trichomoniasis, all birds received Cankerex (MedPet, Newport, WA, USA) in their only drinking water sources at 1 g/L for their first five days in captivity. Following this treatment, to minimize risk of coccidiosis, birds were provided with Endocox in their drinking water at 1.32g/L (L; 2.5% 99 toltrazuril, Jedds Bird Supplies, Anaheim, California, USA) for three consecutive days each week for four weeks. The other four days of each of these weeks, birds were provided with 1 g/L probiotics in their drinking water (Bene-Bac Plus bird and reptile supplement, Pet Ag, Inc., Hampshire, IL USA). After four weeks, Endocox was provided as above for three consecutive days every other week. Probiotics were provided twice weekly in doses as above.

### Responses to initial inoculation

We investigated factors predicting house finch susceptibility, tolerance, eye pathology, and resistance to initial inoculation with *Mycoplasma gallisepticum* (MG) by using an information theoretic model selection approach. This dataset is restricted to finches inoculated with an initial dose of 750 CCU of MG (n=116). We used the same criteria as described in the methods of the main text to assess infection status, initial loads (i.e., resistance), initial tolerance, and maximum initial eye score.

For each analysis, we generated a list of candidate linear models, generalized linear models, or ordinal regressions as appropriate and compared them by AICc by using the 'aictab' function (*bbmle* package; [47]). Predictors included host sex, population, and the interaction between host sex and population. Models <2 ΔAICc for each analysis are reported. In cases where the null has the highest weight, no other models are reported. In cases where there are multiple models <2 ΔAICc and the top model (i.e., highest weighted) is also the most parsimonious, only the top model is reported [48,49]. In cases where there are multiple models <2 ΔAICc, and lower weighted models are nested in the top model, only the top model is reported. For top models, we assessed residuals using the *DHARMa* package [50]. All analyses were carried out in R version 2024.04.2+764 (R Core Team 2024).

*A) Model comparison: Susceptibility to initial inoculation*

Among all finches (n=116), none of our predictors (sex, host population, and the interaction of sex and population) improved predictions of susceptibility to an initial MG inoculation, compared to a null model. See Table S8 for details.

*B) Model comparison: Resistance to initial inoculation*

Among initially infected finches (n = 62), those from the MG-endemic population (VA) had lower loads (i.e., were more resistant) than those from the MG-naive population (AZ; Est = -0.40 ± 0.25). It should be noted that the null model was ranked in the top model set; see Table S9. However, a Kruskal-Wallis rank sum test detected a difference in load among groups categorized by population and sex ($X^2_3$ = 11.04, p = 0.01); see Figure S1. A Dunn test with a Benjamini-Hochberg correction showed this difference existed between female finches from MG-naive and MG-endemic populations, respectively (Z = 3.25, p (adj) = 0.007). This suggests females from the MG-endemic population are more resistant to an initial MG inoculation than females from the MG-naive population.

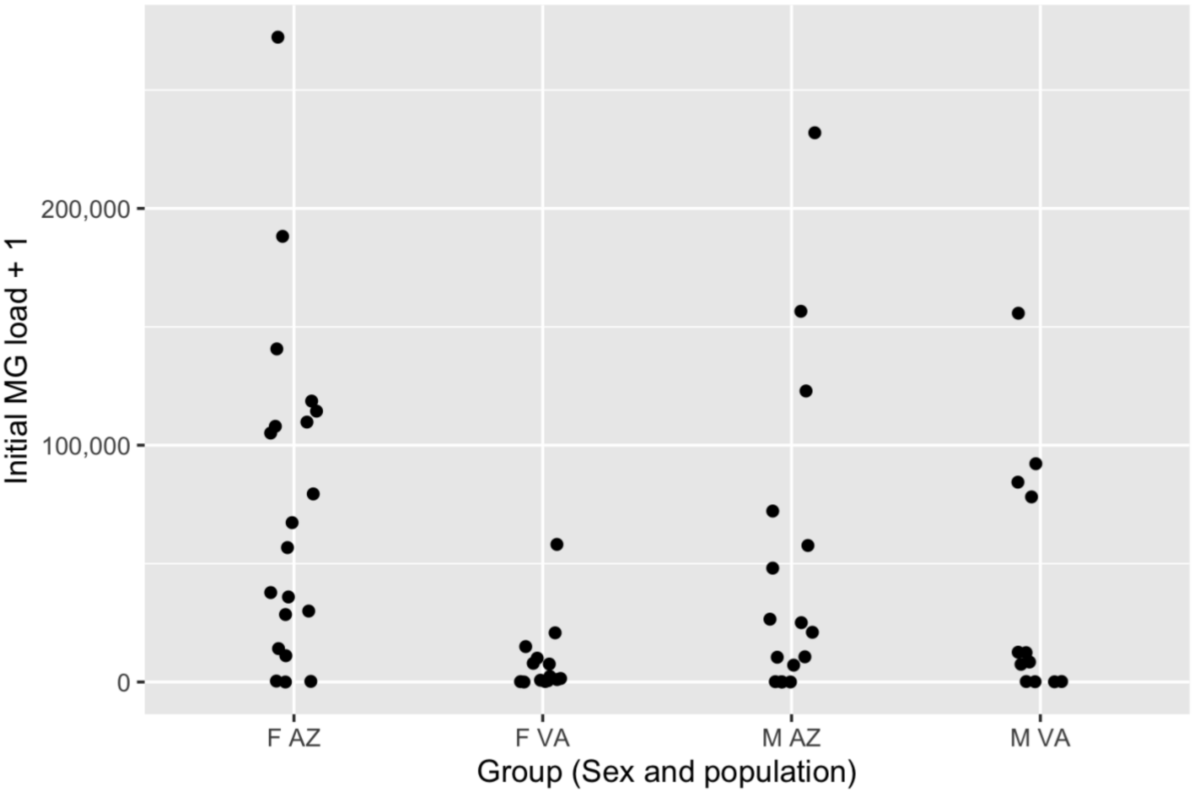


Figure S1. House finch responses to an initial *Mycoplasma gallisepticum* (MG) inoculation. MG loads were assessed 7 days post-inoculation (dppi). Points reflect data from individual birds that were successfully infected; points are jittered for ease of viewing. Data are grouped by host sex (f = female, m = male) and population of origin (az = Arizona, va = Virginia). Y axis is log10 transformed.

*C) Model comparison: Eye pathology following initial inoculation*

Among finches susceptible to an initial MG inoculation (n = 62), initial eye scores were positively associated with initial load (Est = 0.99 ± 0.22), and were lower in finches from the MG-endemic population (VA) than those from the MG-naive population (AZ; Est = -1.96 ± 0.54); see Figure S2 and Table S10.

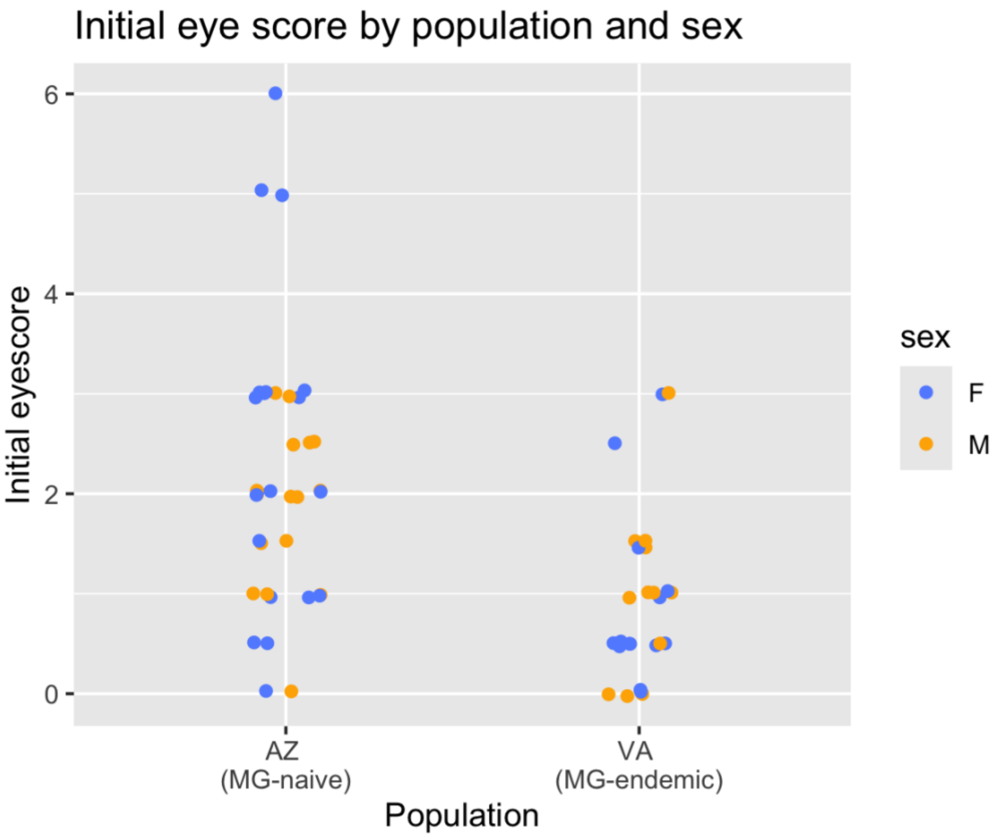


Figure S2. Eye pathology resulting from experimental *Mycoplasma gallisepticum* infection varies by house finch host population. ‘va’ indicates finches from Virginia, a relatively MG-endemic population, while ‘az’ indicates finches from Arizona, a relatively MG-naive population. Pathology is scored on a three point scale for each eye and scores for both eyes are combined; the maximum combined scores for individual finches during the first two weeks following inoculation are shown. Data are jittered for easier viewing. Blue represents females and yellow points represent data from individual males.

*D) Model comparison: Tolerance to initial inoculation*

Among finches susceptible to an initial MG inoculation (n = 62), finches from the MG-endemic population (VA) showed higher tolerance than those from the MG-naive population (AZ; Est = 0.12 ± 0.04); see Figure S3 and Table S11.

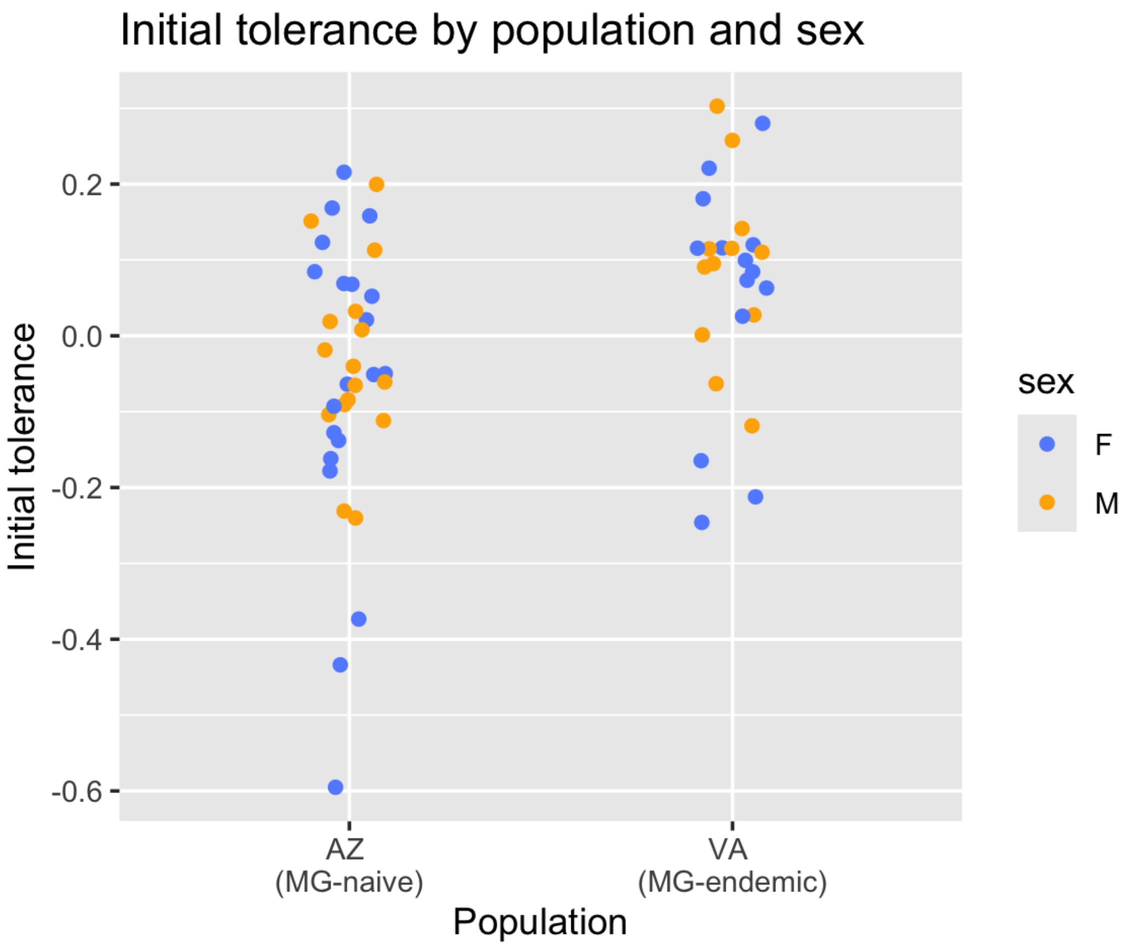


Figure S3. House finches from Virginia (‘VA’) show higher tolerance to an initial *Mycoplasma gallisepticum* inoculation than finches from Arizona (‘AZ’). Shown are tolerance values for finches that were successfully infected following a single MG inoculation (n = 62). Points represent data for individual birds (blue = female) and yellow = male) and are jittered for easier viewing. See methods for further details regarding tolerance quantification.

### Model lists

*A) Susceptibility to the second inoculation (all finches)*

secinf.null: susceptibility ~ 1
secinf.a: susceptibility ~ second dose
secinf.b: susceptibility ~ second dose + population
secinf.c: susceptibility ~ second dose*population
secinf.d: susceptibility ~ second dose + sex
secinf.e: susceptibility ~ second dose*sex
secinf.f: susceptibility ~ second dose + initial susceptibility
secinf.g: susceptibility ~ second dose*initial susceptibility
secinf.h: susceptibility ~ second dose + initial resistance
secinf.i: susceptibility ~ second dose*initial resistance
secinf.j: susceptibility ~ second dose + initial eye score
secinf.k: susceptibility ~ second dose*initial eye score
secinf.l: susceptibility ~ second dose + population+sex
secinf.m: susceptibility ~ second dose + population*sex
secinf.n: susceptibility ~ second dose + initial susceptibility*population
secinf.o: susceptibility ~ second dose + initial susceptibility + population
secinf.p: susceptibility ~ second dose + initial susceptibility*sex
secinf.q: susceptibility ~ second dose + initial susceptibility + sex
secinf.r: susceptibility ~ second dose + initial eye score*sex
secinf.s: susceptibility ~ second dose + initial eye score + sex
secinf.t: susceptibility ~ second dose + initial eye score*population
secinf.u: susceptibility ~ second dose + initial eye score + population
secinf.v: susceptibility ~ second dose + initial resistance*population
secinf.w: susceptibility ~ second dose + initial resistance + population
secinf.x: susceptibility ~ second dose + initial resistance*sex
secinf.y: susceptibility ~ second dose + initial resistance + sex
secinf.z: susceptibility ~ second dose + initial susceptibility:initial tolerance
secinf.aa: susceptibility ~ second dose + initial susceptibility:initial tolerance + sex
secinf.bb: susceptibility ~ second dose + initial susceptibility:initial tolerance + population
secinf.full: susceptibility ~ second dose + sex+population + initial susceptibility + initial resistance + initial eye score + initial susceptibility:initial tolerance

*B) Susceptibility to the second inoculation (in finches susceptible to initial infection)*

secinf2.null: susceptibility ~ 1
secinf2.a: susceptibility ~ second dose
secinf2.b: susceptibility ~ second dose + population
secinf2.c: susceptibility ~ second dose*population
secinf2.d: susceptibility ~ second dose + sex
secinf2.e: susceptibility ~ second dose*sex
secinf2.f: susceptibility ~ second dose + initial resistance
secinf2.g: susceptibility ~ second dose*initial resistance
secinf2.h: susceptibility ~ second dose + initial eye score
secinf2.i: susceptibility ~ second dose*initial eye score
secinf2.j: susceptibility ~ second dose + population + sex
secinf2.k: susceptibility ~ second dose + population*sex
secinf2.l: susceptibility ~ second dose + initial eye score*sex
secinf2.m: susceptibility ~ second dose + initial eye score+sex
secinf2.n: susceptibility ~ second dose + initial eye score*population
secinf2.o: susceptibility ~ second dose + initial eye score + population
secinf2.p: susceptibility ~ second dose + initial resistance*population
secinf2.q: susceptibility ~ second dose + initial resistance + population
secinf2.r: susceptibility ~ second dose + initial resistance*sex
secinf2.s: susceptibility ~ second dose + initial resistance + sex
secinf2.t: susceptibility ~ second dose + initial tolerance
secinf2.u: susceptibility ~ second dose + initial tolerance + population
secinf2.v: susceptibility ~ second dose + initial tolerance + sex
secinf2.w: susceptibility ~ second dose + initial tolerance + sex + population

*C) Resistance to the second inoculation (in all finches susceptible to the second inoculation)*

secresist.null: resistance ~1
secresist.a: resistance ~ second dose
secresist.b: resistance ~ population
secresist.c: resistance ~ sex
secresist.d: resistance ~ initial susceptibility
secresist.e: resistance ~ initial resistance
secresist.f: resistance ~ initial eye score
secresist.g: resistance ~ second dose + population
secresist.h: resistance ~ second dose + sex
secresist.i: resistance ~ initial eye score + population
secresist.j: resistance ~ second dose + initial susceptibility
secresist.k: resistance ~ second dose + initial resistance
secresist.l: resistance ~ second dose + initial eye score
secresist.m: resistance ~ initial susceptibility + population
secresist.n: resistance ~ initial susceptibility + sex
secresist.o: resistance ~ initial resistance + population
secresist.p: resistance ~ initial resistance + sex
secresist.q: resistance ~ initial eye score + sex
secresist.r: resistance ~ sex+population + second dose + initial resistance
secresist.s: resistance ~ sex + population + second dose + initial eye score
secresist.t: resistance ~ sex + population + second dose + initial susceptibility
secresist.u: resistance ~ initial susceptibility:initial tolerance

*D) Resistance to the second inoculation (in AZ finches susceptible to the second inoculation)*

secresist.null2: resistance ~ 1
secresist.a2: resistance ~ second dose
secresist.b2: resistance ~ sex
secresist.c2: resistance ~ initial susceptibility
secresist.d2: resistance ~ initial resistance
secresist.e2: resistance ~ initial eye score
secresist.f2: resistance ~ initial susceptibility:initial tolerance

*E) Tolerance of the second inoculation (in all finches susceptible to the second inoculation)*

sectol.null: tolerance ~ 1
sectol.a: tolerance ~ second dose
sectol.b: tolerance ~ population
sectol.c: tolerance ~ sex
sectol.d: tolerance ~ initial susceptibility
sectol.e: tolerance ~ initial resistance
sectol.f: tolerance ~ initial eye score
sectol.g: tolerance ~ second dose + population
sectol.h: tolerance ~ second dose + sex
sectol.i: tolerance ~ initial eye score + population
sectol.j: tolerance ~ second dose + initial susceptibility
sectol.k: tolerance ~ second dose + initial resistance
sectol.l: tolerance ~ second dose + initial eye score
sectol.m: tolerance ~ initial susceptibility + population
sectol.n: tolerance ~ initial susceptibility + sex
sectol.o: tolerance ~ initial resistance + population
sectol.p: tolerance ~ initial resistance + sex
sectol.q: tolerance ~ initial eye score + sex
sectol.r: tolerance ~ sex + population + second dose + initial resistance
sectol.s: tolerance ~ sex + population + second dose + initial eye score
sectol.t: tolerance ~ sex + population + second dose + initial susceptibility
sectol.u: tolerance ~ initial susceptibility:initial tolerance

*F) Tolerance of the second inoculation (in AZ finches susceptible to the second inoculation)*

sectol2.null: tolerance ~ 1
sectol2.a: tolerance ~ second dose
sectol2.b: tolerance ~ sex
sectol2.c: tolerance ~ initial susceptibility
sectol2.d: tolerance ~ initial resistance
sectol2.e: tolerance ~ initial eye score
sectol2.f: tolerance ~ second dose + sex
sectol2.g: tolerance ~ second dose + initial susceptibility
sectol2.h: tolerance ~ second dose + initial resistance
sectol2.i: tolerance ~ second dose + initial eye score
sectol2.j: tolerance ~ initial susceptibility + sex
sectol2.k: tolerance ~ initial resistance + sex
sectol2.l: tolerance ~ initial eye score + sex
sectol2.m: tolerance ~ initial susceptibility:initial tolerance

*G) Susceptibility to the initial inoculation (all finches)*

priminf.null: susceptibility ~1
priminf.a: susceptibility ~ population
priminf.b: susceptibility ~ sex
priminf.c: susceptibility ~ population + sex
priminf.d: susceptibility ~ population*sex

*<u>H) Resistance to the initial inoculation (all finches susceptible to initial inoculation)</u>*

primresist.null: resistance ~ 1
primresist.a: resistance ~ population
primresist.b: resistance ~ sex
primresist.c: resistance ~ sex + population
primresist.d: resistance ~ sex*population

*I) Initial inoculation pathology (all finches susceptible to initial inoculation)*

glmo0: eye score ~ 1
glmo1: eye score ~ resistance
glmo2: eye score ~ resistance + population
glmo3: eye score ~ resistance*population
glmo4: eye score ~ resistance + sex
glmo5: eye score ~ resistance * sex
glmo6: eye score ~ resistance + sex + population
glmo7: eye score ~ resistance + sex*population
glmo8: eye score ~ resistance*sex + population

*J) Initial inoculation tolerance (all finches susceptible to initial inoculation)*

primtol.null: tolerance ~ 1
primtol.a: tolerance ~ population
primtol.b: tolerance ~ sex
primtol.c: tolerance ~ population*sex
primtol.d: tolerance ~ population + sex

**Tables**

**Table S1.** Sample sizes of house finches (*Haemorhous mexicanus*) given two *Mycoplasma gallisepticum* (MG) inoculations, by host population of origin (AZ = Arizona, VA = Virginia), second dose concentration, and host sex (n = 84).

| Host population | Second MG dose (CCU/mL) | Host sex | n |
|---|---|---|---|
| AZ | 30 | F | 5 |
| AZ | 30 | M | 6 |
| AZ | 100 | F | 5 |
| AZ | 100 | M | 5 |
| AZ | 300 | F | 3 |
| AZ | 300 | M | 6 |
| AZ | 7000 | F | 7 |
| AZ | 7000 | M | 4 |
| VA | 30 | F | 5 |
| VA | 30 | M | 6 |
| VA | 100 | F | 6 |
| VA | 100 | M | 5 |
| VA | 300 | F | 5 |
| VA | 300 | M | 6 |
| VA | 7000 | F | 4 |
| VA | 7000 | M | 6 |

**Table S2.** Generalized linear models predicting house finch (*Haemorhous mexicanus*; n = 84) susceptibility to a second inoculation of *Mycoplasma gallisepticum* bacteria. All models used a binomial error distribution. Shown are parameter estimates (± standard error) for all models <2 ΔAIC. See Model List A in the Supplement for all model details.

| Model | Intercept | Second dose | Initial tolerance | Sex (male) | k | AICc | ΔAICc | Weight |
|---|---|---|---|---|---|---|---|---|
| secinf.aa | -2.66 | $3.90^{-4} \pm 9.88^{-5}$ | -11.14 ± 5.39 | 1.17 ± 0.69 | 4 | 74.23 | 0.00 | 0.30 |
| secinf.z | -1.87 | $3.57^{-4} \pm 9.10^{-5}$ | -11.19 ± 5.52 | | 3 | 75.21 | 0.99 | 0.18 |
| Null | -1.02 | | | | 1 | 98.04 | 23.82 | 0.00 |

**Table S3.** Generalized linear models predicting house finch (*Haemorhous mexicanus*) susceptibility to a second inoculation of *Mycoplasma gallisepticum* (MG) bacteria. Dataset is restricted to finches successfully infected following an initial MG inoculation (n = 37). All models used a binomial error distribution. Shown are parameter estimates (± standard error) for all models <2 ΔAIC. See Model List B in the Supplement for all model details.

| Model | Intercept | Second dose | Initial tolerance | Sex (male) | Population (VA) | k | AICc | ΔAICc | Weight |
|---|---|---|---|---|---|---|---|---|---|
| secinf2.w | -3.70 | $5.11^{-4}$ ± $2.50^{-4}$ | -12.04 ± 6.55 | 3.61 ± 1.89 | -2.62 ± 1.39 | 5 | 32.66 | 0.00 | 0.34 |
| secinf2.j | -3.01 | $5.07^{-4}$ ± $2.09^{-4}$ | | 2.83 ± 1.48 | -2.76 ± 1.28 | 4 | 34.48 | 1.82 | 0.14 |
| secinf2.v | -3.32 | $3.40^{-4}$ ± $1.87^{-4}$ | -12.38 ± 5.63 | 2.25 ± 1.40 | | 4 | 34.57 | 1.91 | 0.13 |
| Null | -1.10 | | | | | 1 | 42.61 | 9.95 | 0.00 |

**Table S4.** Linear models predicting house finch (*Haemorhous mexicanus*) resistance to a second inoculation of *Mycoplasma gallisepticum* (MG) bacteria. Resistance is quantified as MG load at seven days following the second inoculation (lower loads indicate higher resistance). Dataset is restricted to finches successfully infected following a second MG inoculation (n = 22). Shown are parameter estimates (± standard error) for all models <2 ΔAIC. See Model List C in the Supplement for all model details.

| Model | Intercept | Second dose | Initial susceptibility (uninfected) | k | AICc | ΔAICc | Weight |
|---|---|---|---|---|---|---|---|
| secresist.j | 0.89 | $1.73^{-4} \pm 6.99^{-5}$ | 3.01 ± 0.47 | 4 | 72.44 | 0.00 | 0.61 |
| Null | 3.45 | | | 2 | 96.20 | 23.76 | 0.00 |

**Table S5.** Linear models predicting house finch (*Haemorhous mexicanus*) resistance to a second inoculation of *Mycoplasma gallisepticum* (MG) bacteria. Resistance is quantified as MG load at seven days following the second inoculation (lower loads indicate higher resistance). Dataset is restricted to finches from Arizona that were successfully infected following a second MG inoculation (n = 13). Shown are parameter estimates (± standard error) for all models <2 ΔAIC. See Model List D in the Supplement for all model details.

| Model | Intercept | Initial susceptibility (uninfected) | Initial load | Initial eye score | k | AICc | ΔAICc | Weight |
|---|---|---|---|---|---|---|---|---|
| secresist.c2 | 2.02 | 2.75 ± 0.74 | | | 3 | 50.88 | 0.00 | 0.41 |
| secresist.d2 | 4.31 | | -0.64 ± 0.17 | | 3 | 51.02 | 0.13 | 0.38 |
| secresist.e2 | 4.29 | | | -1.09 ± 0.33 | 3 | 52.49 | 1.61 | 0.18 |
| Null | 3.29 | | | | 2 | 57.96 | 7.08 | 0.01 |

**Table S6.** Linear models predicting house finch (*Haemorhous mexicanus*) tolerance to a second inoculation of *Mycoplasma gallisepticum* (MG) bacteria. Tolerance is quantified as residuals of a generalized linear model (binomial family) using MG load at seven days post-inoculation to predict maximum eye score recorded over the first two weeks following inoculation. Analysis is restricted to data from finches successfully infected following a second inoculation (n = 22). Residuals of this model, multiplied by -1, were used as tolerance values; thus, larger values indicate higher tolerance (i.e., relatively less pathology per unit pathogen load relative to the rest of the population). Shown are parameter estimates (± standard error) for all models <2 ΔAIC. See Model List E in the Supplement for all model details.

| Model | Intercept | Second dose | Population (VA) | Initially infected: initial tolerance | Initial susceptibility (infected) | k | AICc | ΔAICc | Weight |
|---|---|---|---|---|---|---|---|---|---|
| Null | $2.22^{-13}$ | | | | | 2 | 4.57 | 0.00 | 0.19 |
| sectol.a | $-7.83^{-2}$ | $1.74^{-5} \pm 1.59^{-5}$ | | | | 3 | 6.00 | 1.44 | 0.09 |
| sectol.b | -0.04 | | 0.10 ± 0.11 | | | 3 | 6.34 | 1.77 | 0.08 |
| sectol.u | -0.02 | | | -1.09 ± 1.19 | | 3 | 6.36 | 1.79 | 0.08 |
| sectol.d | 0.06 | | | | 0.10 ± 0.11 | 3 | 6.40 | 1.83 | 0.08 |

**Table S7.** Linear models predicting house finch (*Haemorhous mexicanus*) tolerance to a second inoculation of *Mycoplasma gallisepticum* (MG) bacteria. Tolerance is quantified as residuals of a generalized linear model (binomial family) using MG load at seven days post-inoculation to predict maximum eye score recorded over the first two weeks following inoculation. Residuals of this model, multiplied by -1, were used as tolerance values; thus, larger values indicate higher tolerance (i.e., relatively less pathology per unit pathogen load relative to the rest of the population). Dataset is restricted to finches from Arizona that were successfully infected following a second MG inoculation (n = 13). Shown are parameter estimates (± standard error) for all models <2 ΔAIC. See Model List E in the Supplement for all model details.

| Model | Intercept | Initial susceptibility (uninfected) | Second dose | k | AICc | ΔAICc | Weight |
|---|---|---|---|---|---|---|---|
| sectol2.g | $-5.58^{-2}$ | -0.33 ± -0.12 | $3.86^{-5} \pm 1.82^{-5}$ | 4 | 6.79 | 0.00 | 0.29 |
| sectol2.c | 0.10 | -0.33 ± 0.12 | | 3 | 7.29 | 0.50 | 0.23 |
| Null | -0.04 | | | 2 | 8.53 | 1.74 | 0.12 |

**Table S8.** Generalized linear models predicting house finch (*Haemorhous mexicanus*; n = 116) susceptibility to an initial inoculation of *Mycoplasma gallisepticum* (MG) bacteria. All models use a binomial family. Shown are parameter estimates (± standard error) for all models <2 ΔAIC. See Model List G in the Supplement for all model details.

| Model | Intercept | Sex (M) | Population (VA) | k | AICc | ΔAICc | Weight |
|---|---|---|---|---|---|---|---|
| Null | 0.14 | | | 1 | 162.29 | 0.00 | 0.43 |
| priminf.b | 0.31 | -0.34 ± 0.38 | | 2 | 163.52 | 1.23 | 0.24 |
| priminf.a | 0.25 | | -0.25 ± 0.37 | 2 | 163.91 | 1.62 | 0.19 |

**Table S9.** Generalized linear models predicting house finch (*Haemorhous mexicanus*) resistance to an initial inoculation of *Mycoplasma gallisepticum* (MG) bacteria. Resistance is quantified as MG load at seven days following the second inoculation (lower loads indicate higher resistance). All models use a negative binomial family. Data are restricted to finches that were successfully infected following initial inoculation (n = 62). Shown are parameter estimates (± standard error) for all models <2 ΔAIC. See Model List H in the Supplement for all model details.

| **Model** | **Intercept** | **Population (VA)** | **Sex (M)** | **Sex (M) * population (VA)** | **k** | **AICc** | **ΔAICc** | **Weight** |
|---|---|---|---|---|---|---|---|---|
| primresist.a | 10.90 | -0.40 ± 0.25 | | | 3 | 1378.41 | 0.00 | 0.33 |
| Null | 10.75 | | | | 2 | 1378.81 | 0.40 | 0.27 |
| primresist.d | 11.10 | | | 0.81 ± 0.50 | 5 | 1379.93 | 1.52 | 0.16 |
| primresist.c | 10.98 | -0.43 ± 0.25 | -0.17 ± 0.25 | | 4 | 1380.22 | 1.81 | 0.14 |

**Table S10.** Ordinal regression models predicting house finch (*Haemorhous mexicanus*) pathology following an initial inoculation of *Mycoplasma gallisepticum* (MG) bacteria. Maximum initial eye score is the highest eye score recorded within the first two weeks following initial inoculation. Data are restricted to finches that were successfully infected following initial inoculation (n = 62). Shown are parameter estimates (± standard error) for all models <2 ΔAIC. Full models and parameters available in code. See Model List I in the Supplement for all model details.

| Model | Population (VA) | Initial load | Initial load * population (VA) | k | AICc | ΔAICc | Weight |
|---|---|---|---|---|---|---|---|
| glmo2 | -1.96 ± 0.54 | 0.98 ± 0.21 | | 10 | 231.16 | 0.00 | 0.50 |
| glmo3 | | | -0.18 ± 0.42 | 11 | 233.07 | 1.90 | 0.19 |
| Null | | | | 9 | 268.39 | 37.22 | 0.00 |

**Table S11.** Linear models predicting house finch (*Haemorhous mexicanus*) tolerance to an initial inoculation of *Mycoplasma gallisepticum* (MG) bacteria. Tolerance is quantified as residuals of a generalized linear model (binomial family) using MG load at seven days post-inoculation to predict maximum eye score recorded over the first two weeks following inoculation. Residuals of this model, multiplied by -1, were used as tolerance values; thus, larger values indicate higher tolerance (i.e., relatively less pathology per unit pathogen load relative to the rest of the population). Data are restricted to finches that were successfully infected following initial inoculation (n = 62). Shown are parameter estimates (± standard error) for all models <2 ΔAIC. See Model List J in the Supplement for all model details.

| Model | Intercept | Population (VA) | Sex (M) | k | AICc | ΔAICc | Weight |
|---|---|---|---|---|---|---|---|
| primtol.a | -0.05 | 0.12 ± 0.04 | | 3 | -46.12 | 0.00 | 0.60 |
| primtol.d | -0.07 | 0.12 ± 0.04 | 0.03 ± 0.04 | 4 | -44.54 | 1.59 | 0.27 |
| Null | $2.68^{-15}$ | | | 2 | -40.07 | 6.06 | 0.03 |
